# Local thalamic interneurons drive spindle termination and enable sleep-dependent learning

**DOI:** 10.1101/2025.09.26.678870

**Authors:** Jane Simko, Tamina Keira Ramirez, Caryn Martin, Sally Leung, Eleni A. Sung, Christopher D. Makinson

## Abstract

The thalamus is central to fundamental brain functions including sensation, attention, and sleep through the precise generation and regulation of neuronal ensemble oscillatory activity. Sensory thalamic circuits are considered feedforward structures, lacking lateral connectivity, while recurrence in the network is mediated by interactions with inhibitory neurons of the thalamic reticular nucleus. Here, we define previously uncharacterized functional roles of local thalamic interneurons, a component of the sensory thalamus whose function has remained unexplored. We demonstrate that local interneuron activation induces rebound oscillations in thalamocortical relay neurons *ex vivo* and neocortical spindles *in vivo*, while their inhibition increases spindle occurrence, overall spindle duration and impairs sensory learning. Our findings reveal that local thalamic interneurons have shared and complementary functions to those of thalamic reticular neurons and are required for proper spindle formation and sleep-dependent learning. Together, this work establishes an important neural substrate of thalamocortical circuit function.

## Introduction

The ventral posterior (VP) somatosensory thalamus plays a central role in fundamental brain processes including sleep and sensation [1] [2] [3] [4] [5] [6]. Importantly, local interneurons in the somatosensory thalamus represent an evolutionarily conserved cell type broadly present across mammalian species, including humans where they represent nearly half of the neuronal population [7]. However, local interneurons in the mouse somatosensory thalamus have only recently been characterized after having been overlooked due to their contrasting sparsity (approximately 1-5%) [7] [8] [9]. Though somatically sparse in rodents, ventral posterior local interneurons (VPi) are morphologically expansive with arborizations that encompass entire thalamic subnuclei [9] enabling each of them to establish contacts with hundreds of synaptic partners and influence widespread network activity.

Oscillatory events, particularly spindles during slow-wave sleep, are thought to be generated through interactions between the deep pyramidal layers of the neocortex, the excitatory VP thalamocortical (TC) relays, and the neighboring thalamic reticular nucleus (TRN) [1] [2] [3] [4] [10]. Both TC and TRN neurons exhibit bursting in response to reciprocal synaptic signaling [11] [12] [13] [14] [15], creating a positive feedback loop that generates rhythmic activity. These rhythms engage the larger thalamocortical loop in events such as sleep spindles where they appear in electroencephalogram (EEG) recordings as 9-16 Hz activity in adult mice [3] [16].

Sleep spindle regulation has been studied within a framework largely confined to TC, TRN, and cortical neuron interactions. Neocortical spindles during NREM sleep have been shown to be initiated following phasic optogenetic activation of the TRN, providing direct evidence for their role in switching the firing mode of TC neurons [17]. Layer 6 cortical UP states are a plausible naturalistic stimulus triggering TRN neurons to initiate thalamic rhythms [18] [19]. Once initiated, spindle maintenance and synchrony rely on electrical synapses between TRN neurons, which have been shown to reinforce and sustain the coordinated burst firing that underlies these rhythms [20] [21] [22] [23] [3] [24] [25]. Cortical inputs have also been proposed to enhance global rhythmic synchrony through activation of TRN neurons [26] [27] [28], though rhythmic oscillations in the thalamus can be induced and sustained *ex vivo*, isolated from the cortex, indicating that the thalamus can act as a self-contained rhythm generator [29] [30].

Multiple theories have emerged to explain the mechanisms underlying sleep spindle desynchronization and eventual termination. Among them, TRN↔TRN lateral inhibition has been identified as a key factor in thalamic desynchronization, though the role of intra-TRN synaptic connectivity in the mature thalamus remains controversial as these connections are significantly reduced over the course of development [31] [32] [33] [34] [35] [36] [37] [25] [38].

Research into the precise mechanisms underlying local thalamic spindle regulation is ongoing. However, the stereotyped duration of spindles [3] and the sparsity of TC neuron bursting during spindles [39] [40] [41] suggest the presence of robust inhibitory feedback mechanisms that tightly control synchrony and desynchrony. VPi neurons have not yet been investigated in the context of oscillatory events though recent findings have shown that they also possess the ability to generate bursts of action potentials when activated from a hyperpolarized state [9]. Notably, VPi neurons directly inhibit TC relays [9], though this interaction is not reciprocal, raising the question of whether VPi neurons are solely involved in feedforward inhibition in sensory signal processing or if they also contribute to thalamocortical rhythms.

Here, using a Sox14-Cre mouse line to specifically target VPi neurons, we demonstrated that activation of VPi neurons induces TC rebound oscillations in slice preparations and neocortical spindles in behaving mice while inhibition increases spindle incidence, lengthens overall spindle duration and impairs sleep-dependent learning. While TRN neurons have long been recognized as a key inhibitory source of spindle regulation, this study reveals that VPi neurons work in concert with the TRN, regulating spindle initiation and desynchronization. These results highlight a critical and unique contribution of VPi neurons to thalamocortical rhythms with implications for learning and memory consolidation during slow-wave sleep and disorders of the thalamocortical system such as sleep impairment and epilepsy. The dual roles of VPi neurons, bridging feedforward inhibition and rhythm modulation, provide new insights into how they contribute to the complex orchestration of sensory and sleep-related processes.

## Results

### VPi neurons encode movement-related activity

It has previously been shown through *ex vivo* electrophysiology that VPi neurons receive inputs from the medial lemniscus carrying sensory information from the face and body [9] [42] [43] [44] [45] [46]. To measure VPi neuron activity *in vivo* during movement in wake states, we examined the activity patterns of VPi, TC, and TRN neurons by recording GCaMP8m signals via fiber photometry alongside EEG and EMG over a 24-hour period (**Fig. 1a,b**). To selectively target VPi neurons in the VP, we used a Sox14-Cre mouse line. Validation of Cre expression was performed using a local injection of a Cre-dependent mCherry virus, revealing that mCherry-positive neurons co-localized with Gad67 antibodies (73% of cells expressed both mCherry and Gad67, while 27% expressed only mCherry, **Extended Data Fig. 1a-c,e**). Sox14-positive neurons were also detected in the dorsal lateral geniculate nucleus (LGN), where they were more abundant and also co-localized with Gad67 (72.0% of cells expressed both mCherry and Gad67, while 24.4% expressed only anti-Gad67 and 3.6% expressed only mCherry, **Extended Data Fig. 1a,b, d-e**). To measure TC GCaMP activity, we injected CaMKII-GCaMP8m into Sox14-Cre-positive, PV-Cre-positive, or wildtype mice. For VPi activity, we injected FLEX-GCaMP8m into Sox14-Cre-positive mice, while TRN activity was observed by injecting FLEX-GCaMP8m into PV-Cre-positive mice (**Fig. 1c**). GCaMP8m successfully targeted TC, VPi, and TRN populations, as validated by slice and *in vitro* functional imaging for TC and VPi neurons (**Fig. 1c, Extended Data Fig. 2**). No major differences in general sleep characteristics were observed between the three groups of mice targeted for TC, VPi, and TRN recordings (**Extended Data Fig. 3**).

**Fig. 1.**
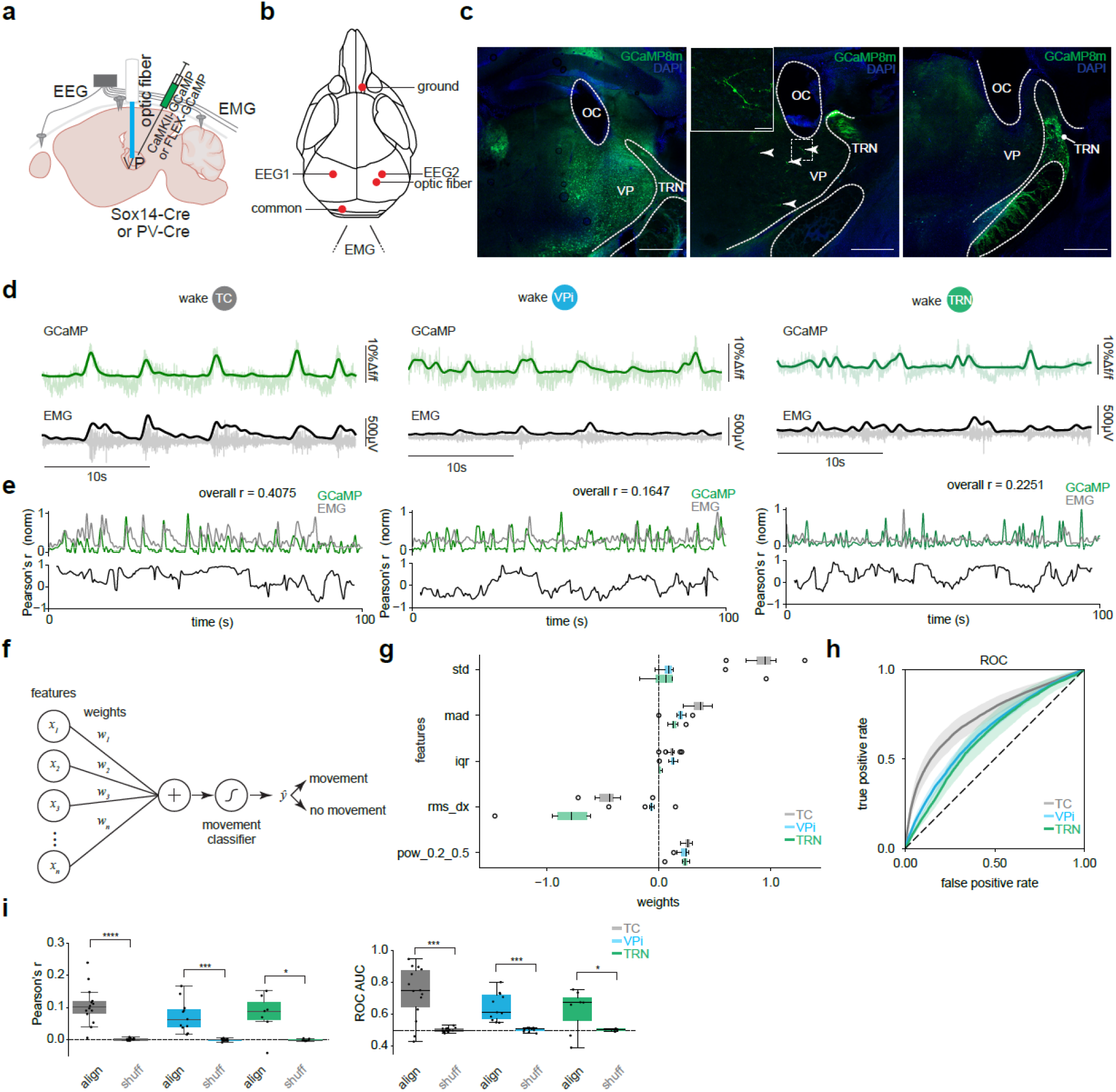
TC VPi and TRN neurons encode movement-related activity. **a**, Stereotaxic injection of either AAV9-CaMKII-GCaMP8m or AAV9-FLEX-GCaMP8m into the VP of Sox14-Cre-positive, PV-Cre-positive, or wildtype mice. **b**, EEG leads are implanted into left and right somatosensory areas. The optic fiber is implanted above the VP thalamus. **c**, Optic fiber cannulae (OC) location in relation to the VP and TRN thalamic nuclei. **d**, Example traces of TC, VPi and TRN GCaMP activity in green and EMG. GCaMP and EMG signals are half-wave rectified and low pass filtered. The resulting EMG signal is multiplied by 10 for visibility. **e**, Example time-locked smoothed GCaMP and EMG signals and a rolling Pearson’s r correlation with a 5 second window. **f**, Schematic of the linear classifier used to predict movement from GCaMP features. Five features were extracted per 10s segment and weighted to classify movement vs. no movement. **g**, Feature weights for each population. **h**, Receiver operating characteristic (ROC) curves of leave-one-mouse-out cross-validation showing above-chance prediction for TC, VPi, and TRN populations. Shaded regions indicate SEM. **i**. Left: 10 randomly sampled GCaMP and EMG time-locked (aligned) segments averaged overall Pearson’s r, compared to 10 shuffled GCaMP and EMG signals. Random sampling iterations occurred 100 times. TC *P* = 6.1035e-05, VPi *P* = 0.0005, TRN *P* = 0.0156, one-sided Wilcoxon signed-rank test. Right: ROC AUC vs. shuffled-label AUC. TC *P* = 0.0004, VPi *P* = 0.0004, TRN *P* = 0.0391, one-sided Wilcoxon signed-rank test. TC *n* = 14 mice, VPi *n* = 11 mice, TRN *n* = 7 mice. *P* < 0.0001 (****), *P* < 0.001 (***), *P* < 0.05 (*). Data represented as mean ± SEM. std: standard deviation, mad: median absolute deviation, iqr: interquartile range, rms_dx: root mean square of successive differences, pow_0.2_0.5: relative power in the 0.2-0.5Hz frequency band.

Two EEG channels (right and left hemisphere somatosensory cortex) and two EMG channels were used to automatically classify behavioral states via a convolutional neural network-based sleep scoring system. To assess movement-related activity, we randomly sampled segments of right somatosensory cortex EEG and corresponding EMG recordings from wake periods. Pearson correlation coefficients between aligned (“align”) EEG and EMG signals were computed and compared to those from shuffled (“shuff”) EEG-EMG segments. Similar to TC and TRN neurons, VPi activity was highly correlated with movement (**Fig. 1d-e,i)**.

To test whether GCaMP signals alone could predict movement, we trained a linear classifier using features derived from the calcium signal, independent of EMG (**Fig. 1f**). Each 10s segment was reduced to only five summary statistics: standard deviation (std), median absolute deviation (mad), interquartile range (iqr), root mean square of successive differences (rms_dx), and relative low-frequency power (pow_0.2_0.5), which were weighted to classify movement vs. no-movement epochs (**Fig. 1g**). Classifiers trained on TC, VPi, and TRN populations all achieved above-chance performance, as shown by receiver operating characteristic (ROC) curves from leave-one-mouse-out cross-validation (**Fig. 1h**). For each population, ROC curves from individual folds were averaged to obtain the mean classifier performance. TC neurons provided the strongest decoding, whereas VPi and TRN populations contributed weaker but still significant predictive power. To establish a baseline, the same analysis was repeated with shuffled ground truth labels. Comparison of aligned vs. shuffled controls further confirmed that the calcium signals carried movement information (**Fig. 1i**). Across all populations, Pearson’s correlation coefficients between GCaMP and EMG were higher for aligned data than shuffled EEG-EMG segments, and aligned ROC area under the curve (AUC) values exceeded those obtained with shuffled ground truth labels.

### VPi neurons participate in sleep spindles

Having established that VPi neurons are functionally active during movement, we next investigated whether they are also active during sleep, particularly during sleep spindles. Using a validated spindle detection algorithm [47] (**Fig. 2a**), we randomly sampled segments containing detected (“det”) sleep spindles, constrained to the 9-16 Hz frequency range. These segments were then compared to shuffled (“shuff”) 2s segments randomly selected from any portion of NREM sleep. GCaMP fiber photometry and EEG signals revealed a clear waxing and waning pattern for each targeted cell population, whereas this activity was absent in the shuffled data (**Fig. 2b**). Additionally, spindle-detected segments exhibited a distinct normalized decibel peak within the 9-16 Hz range, whereas shuffled segments showed only a modest peak, likely reflecting the inclusion of all phases of NREM sleep (**Fig. 2b**).

**Fig. 2.**
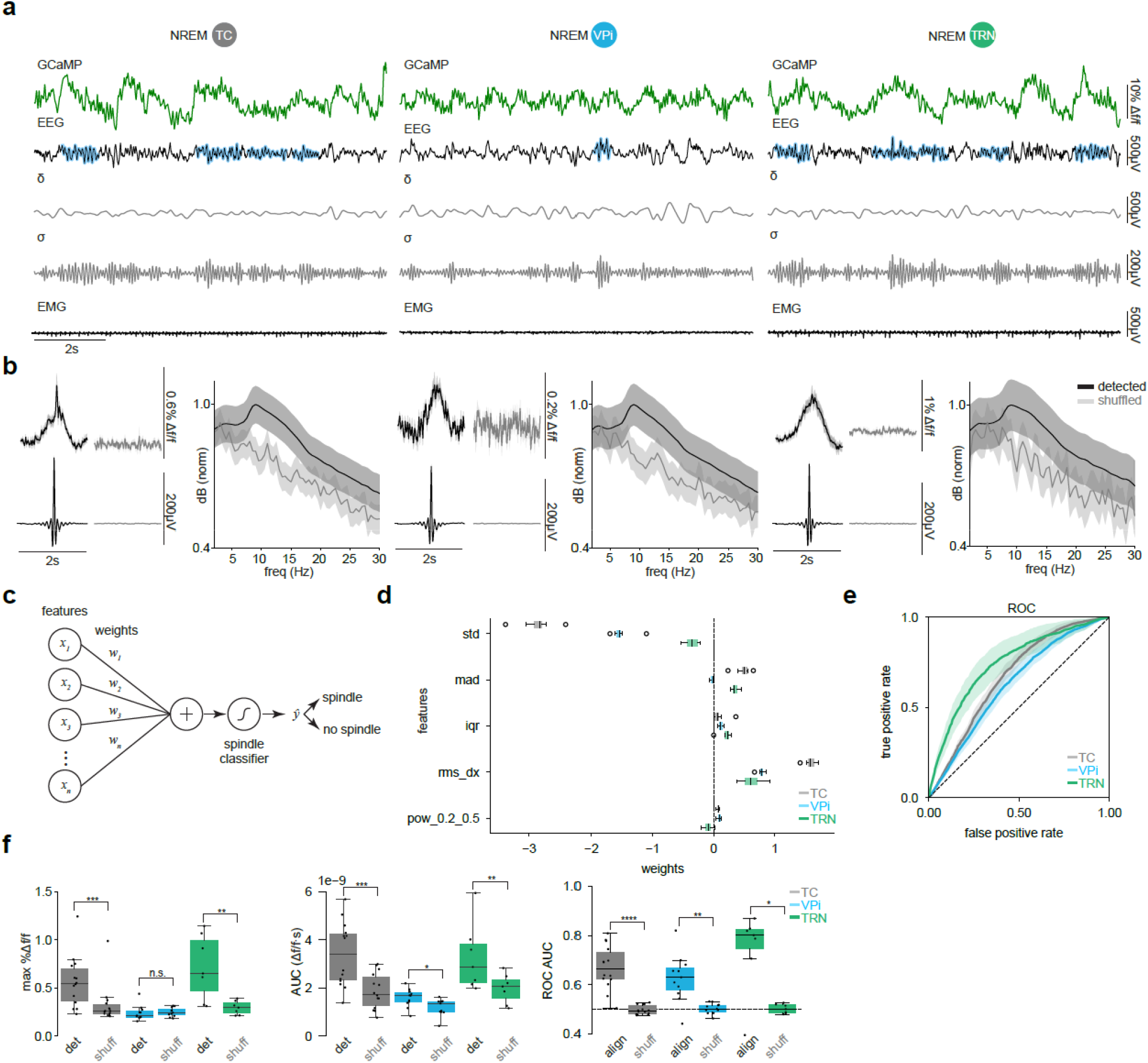
TC VPi and TRN neurons participate in sleep spindles. **a**, Example GCaMP signal, right somatosensory EEG, delta band filtered EEG, sigma band filtered EEG, and EMG. Algorithmically detected sleep spindles shown with a blue mask. **b**, Comparing 100 randomly sampled groups of 2s averaged GCaMP activity and EEG detected spindles vs. 100 shuffled 2s segments of NREM recordings. Random sampling iterations occurred 100 times. EEG or GCaMP segments with mean values exceeding a z-score threshold of 5 were excluded from analysis. Power spectral density plots of detected vs. shuffled 2s segments of EEG data. **c**, Schematic of the linear classifier used to predict spindle presence from GCaMP features. Five features were extracted per 10s segment and weighted to classify spindle vs. no spindle. **d**, Feature weights for each population. **e**, Receiver operating characteristic (ROC) curves of leave-one-mouse-out cross-validation showing above-chance prediction for TC, VPi, and TRN populations. Shaded regions indicate SEM. **f**, Left: Maximum %Δf/f for detected and shuffled events. TC *P* = 0.0006, VPi *P* = 0.8799, TRN *P* = 0.0078 Middle: Area under the curve (AUC) of Δf/f responses for detected and shuffled events. TC *P* = 0.0002, VPi *P* = 0.0210, TRN *P* = 0.0078; Right: ROC AUC vs. shuffled-label AUC. TC *P* = 6.1035e-5, VPi *P* = 0.0015, TRN *P* = 0.0156, one-sided Wilcoxon signed-rank test. TC *n* = 14 mice, VPi *n* = 11 mice, TRN *n* = 7 mice. *P* < 0.0001 (****), *P* < 0.001 (***), *P* < 0.01(**), *P* < 0.05 (*), n.s. = not significant. Data represented as mean ± SEM. std: standard deviation, mad: median absolute deviation, iqr: interquartile range, rms_dx: root mean square of successive differences, pow_0.2_0.5: relative power in the 0.2-0.5Hz frequency band.

Interestingly, VPi, TC, and TRN neurons also displayed a similar pattern of activity during REM sleep, specifically within the mu frequency band (8-12 Hz, **Extended Data Fig. 4**). Notably, GCaMP activity in TC, VPi, and TRN peaked slightly after the averaged center of the EEG waxing and waning cycle (**Extended Data Fig. 4b**). This temporal shift contrasts with sleep spindles, where thalamic activity is highly synchronized with neocortical EEG spindles.

To assess whether GCaMP signals carried information about spindle events, we applied the same linear classification framework as for movement, now trained to distinguish spindle vs. non-spindle epochs using only calcium features, independent of EEG (**Fig. 2c**). Each 10s segment was summarized by five features weighted to classify spindle occurrence (**Fig. 2d**). Using leave-one-mouse-out cross-validation, ROC curves revealed that all three populations supported above-chance decoding, with TRN neurons showing the strongest predictive performance (**Fig. 2e**). Consistent with their established role in spindle generation, TRN GCaMP activity most reliably anticipated spindle events, whereas TC and VPi populations provided weaker but still significant predictive power. The same cross-validation scheme was repeated with shuffled ground-truth labels. Comparison with shuffled data confirmed that the detected spindles were associated with distinct calcium dynamics: max Δf/f values and AUC were significantly elevated in GCaMP signals of detected spindles across all cell populations (**Fig. 2f**). Furthermore, ROC AUC values were consistently higher for aligned (“align”) data compared to that of shuffled (“shuff”) ground-truth labels across all populations, highlighting that GCaMP activity contains predictive information about spindle timing even in the absence of direct EEG input.

### Entrainment of sleep spindles by VPi neuron activation

Given that VPi neurons synapse onto TC neurons [9], we sought to examine how optogenetic stimulation of VPi neurons *ex vivo* affects TC neuron rebound activation (**Fig. 3a,b**). We injected Cre-dependent ChRmine into the VP thalamus of Sox14-Cre mice (**Extended Data Fig. 5, Fig. 3a**) and recorded activity using a 16-channel linear array probe placed across the VP in horizontal slices (**Fig. 3a**). Optostimulation with amber light (595nm) at 1Hz (200ms light-on, 30s total) induced rhythmic activity in putative TC neurons, which occurred more often during light-off periods (**Fig. 3c**). Conversely, in some instances, putative VPi neuron activity was correlated with light-on periods (**Fig. 3c**). Considering the well-established role of TRN neurons in driving TC rebound oscillations, we performed a parallel experiment by expressing ChR2 in PV-positive TRN neurons (**Fig. 3a**). Similar to VPi optostimulation, TRN activation induced rebound bursting in putative TC neurons during light-off periods, while putative TRN neurons were active during light-on periods (**Fig. 3d**). Neurons within the VP were sorted across all slices. Pearson correlation analysis of neuronal activity during the first 30s of optostimulation revealed that putative TC neurons were negatively correlated with stimulation, while putative VPi neurons were positively correlated. In the subsequent 30s without stimulation, these correlations approached zero (**Fig. 3e**). Latency analysis showed that putative TC neurons exhibited a delay of 0.4997 ± 0.0130s compared to the start of light on, whereas putative VPi neurons had a shorter latency of 0.2377 ± 0.0733s (*P* = 0.0179; one-sided Mann-Whitney U test; **Fig. 3e**). Similarly, in TRN optostimulation experiments, putative TC and TRN neurons displayed comparable correlation patterns, with TC neuron latency at 0.3816 ± 0.0369s and TRN neuron latency at 0.1378 ± 0.0356s (*P* = 0.0023; one-sided Mann-Whitney U test; **Fig. 3f**).

**Fig. 3.**
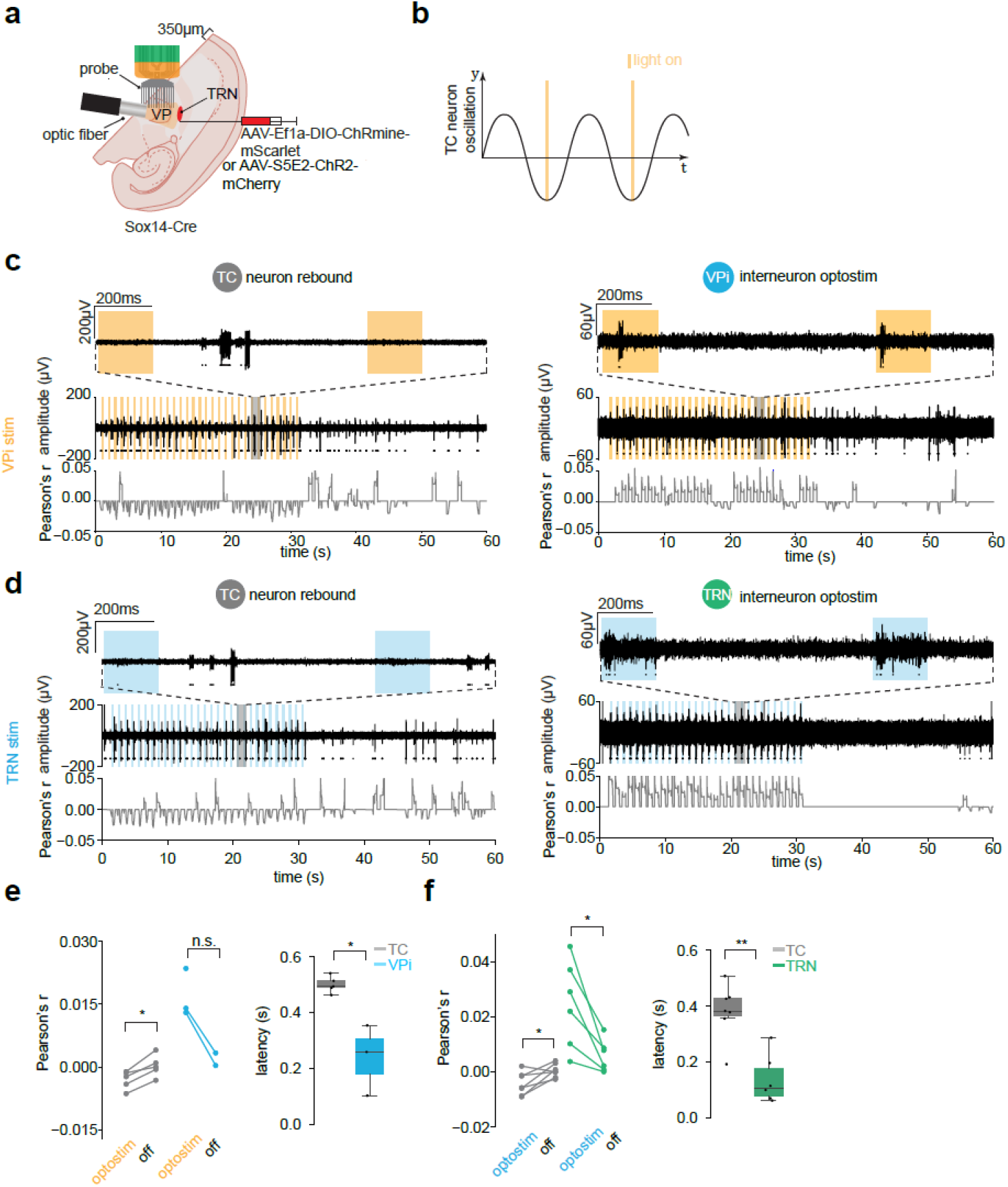
Initiation of TC rebound bursts ex vivo by VPi and TRN neuron activation. **a**, Stereotaxic injection of either AAV5-Ef1a-DIO-ChRmine-oScarlet or AAV9-S5E2-ChR2-mCherry (bilateral) into the VP of Sox14-Cre mice. Experimental setup using a 16 channel NeuroNexus probe and optic fiber emitting 595nm or 465nm light. **b**, Light is turned on for 0.2s at 1Hz. **c**, Left: putative TC neurons rebound burst in light-off periods of VPi optostimulation. Window shaded in grey shown on top axis. Rolling Pearson’s r coefficient shown comparing light-on periods vs. spike detections. Detected neurons shown with black dots (*n* = 2 mice). Right: same as in left panel with putative VPi neurons. VPi neurons are conversely activated during light-on periods. (*n* = 2 mice). **d**, Left: putative TC neurons rebound burst in light-off periods of TRN optostimulation. Window shaded in grey shown on top axis. Rolling Pearson’s r coefficient shown comparing light-on periods vs. spike detections. Detected neuron shown with black dots (*n* = 1 mouse). Right: same as in left panel with putative TRN neurons. TRN neurons are conversely activated during light-on periods. (*n* = 1 mouse). **e**, Left: Comparison of the first 30 seconds of overall Pearson’s r with the last 30 seconds for TC and VPi neurons. TC *P* = 0.0312, VPi n.s., one-sided Wilcoxon signed-rank test. Right: Boxplot of latency values for all detected neurons. *P* = 0.0179, one-sided Mann-Whitney U test. TC *n* = 5, VPi *n* = 3 cells of at least 30s duration, *n* = 2 cells of 60s duration). **f**, same as in E for TRN stim experiments. TC *P* = 0.0234, TRN *P* = 0.0156, one-sided Wilcoxon signed-rank test, *P* = 0.0023, one-sided Mann-Whitney U test. TC *n* = 7 cells, TRN *n* = 6 cells. *P* < 0.01(**), *P* < 0.05 (*), n.s. = not significant.

To determine whether VPi neuron activation also induces TC rebound bursts *in vivo*, we recorded either TC or VPi calcium activity in Sox14-Cre mice injected with Cre-dependent ChRmine into the VP (**Fig. 4a**). We likewise observed the effects of TRN neuron activation on TC or TRN calcium activity in PV-Cre mice injected with Cre-dependent ChRmine (**Fig. 4a**). The 24hr recording day was separated into thirds such that when a 10 second period of REM was detected by our model in the first third, optostimulation would be delivered for 10 seconds. Wake and NREM were the cues for delivering optostimulation in the second and third portions of the 24hr period respectively. Light was delivered at either 0.5, 1, or 1.5Hz per 24hr recording period with a light on of 200ms (**Fig. 4b**). Predictably, VPi neurons showed a rise of activity following each optostimulation for all wake, NREM, and REM states. During wake and REM states, TC calcium activity was suppressed and antiphase to VPi activity, likely due to VPi inhibition of TC neurons. However, during NREM, TC neurons exhibited a peak of calcium activity following each optostimulation (**Fig. 4c**). Considering that this occurs only during NREM, the ability of VPi neurons to initiate TC rebound bursts may be state-dependent. The corresponding TC and TRN activity from TRN activation followed the same pattern as that of VPi activation and also exhibited a NREM state-dependence for TC rebound bursts (**Fig. 4c**). Optostimulation of mice lacking opsin expression failed to elicit any calcium response (**Fig. 4c**) and a comparison of general sleep characteristics of baseline recordings versus their corresponding VPi-stim or TRN-stim recordings showed no major differences (**Extended Data Fig. 6**) though there were some significant differences in other mouse group comparisons.

**Fig. 4.**
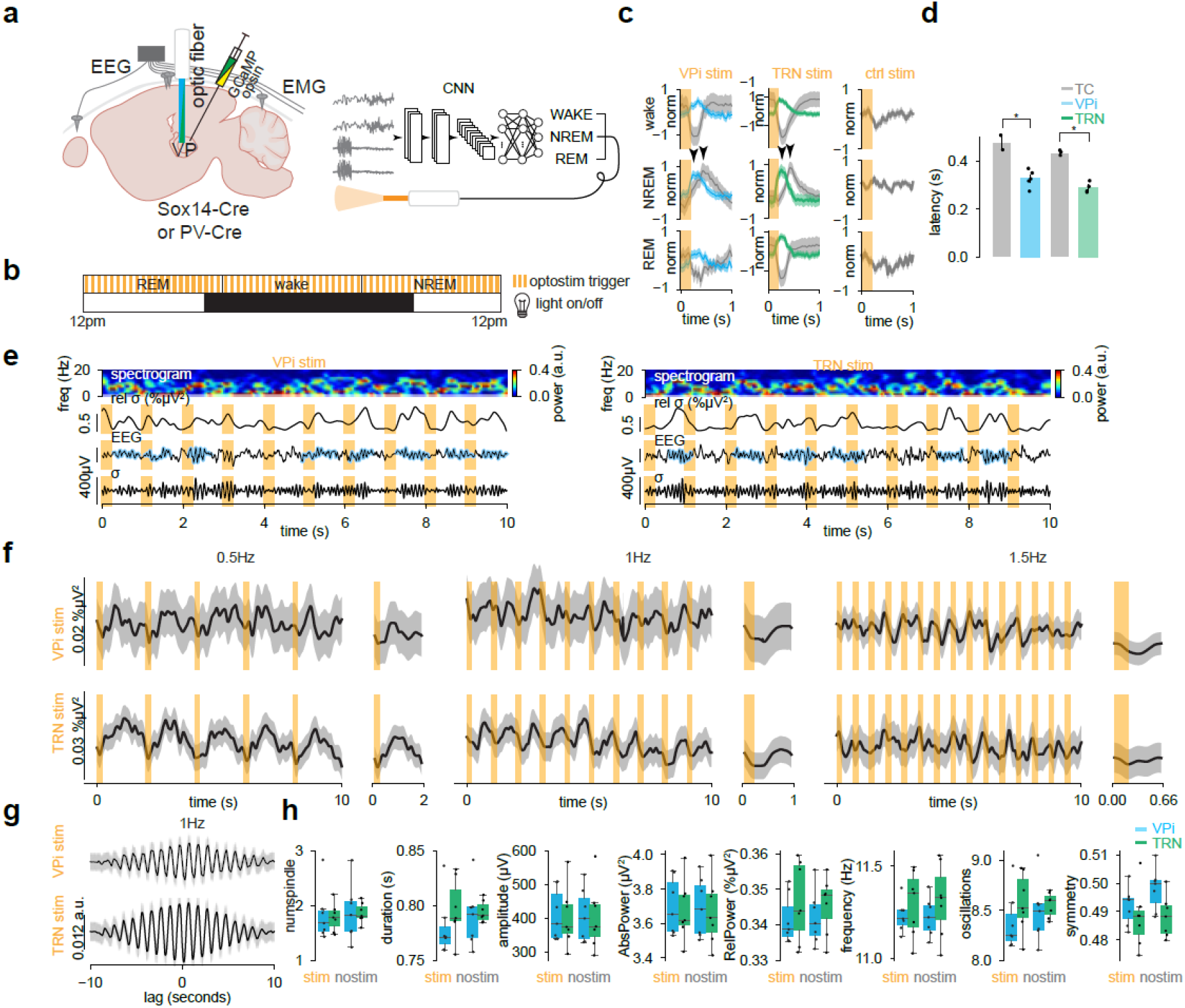
Entrainment of sleep spindles by VPi and TRN neuron activation. **a**, Stereotaxic injection of GCaMP and AAV5-Ef1a-DIO-ChRmine-oScarlet in Sox14-Cre of PV-Cre mice. EEG leads are implanted with screws into left and right somatosensory areas (see Fig. 1). An optic fiber is implanted into the VP. **b**, VPis or TRNs are optostimulated in a closed loop design according to time blocks. 24 hours are split into thirds. REM, wake and NREM events for each third trigger optostimulation periods for 10s. Light is turned on for 0.2s at 0.5, 1, or 1.5Hz. **c**, 100 randomly sampled 10s segments were drawn from NREM and wake. 20 samples of REM were taken due to their sparsity. Random sampling iterations occurred 100 times. Segments with mean values exceeding a z-score threshold of 1 were excluded from analysis. Only 1Hz stimulation recordings shown. TC (*n* = 2 mice) and VPi (*n* = 5 mice) GCaMP responses from VPi optostimulation (left) and TC (*n* = 3 mice) and TRN (*n* = 4 mice) GCaMP responses from TRN optostimulation (middle). GCaMP responses to optostimulation from mice injected without an opsin (*n* = 2 mice; right). **d**, Latency of TC neurons vs VPi or TRN neurons from VPi or TRN optostimulation respectively during NREM 1Hz protocols. Arrows show the approximate peak location. VPi stim: TC latency = 0.4759 ± 0.0302s, VPi latency = 0.3271 ± 0161s, *P* = 0.0476; TRN stim: TC latency = 0.4351 ± 0.0063s, TRN latency = 0.2892 ± 0.0109s, *P* = 0.0286, one-sided Mann-Whitney U test. **e**, Example data from VPi (left) and TRN (right) optostimulation experiments. Right somatosensory EEG data is shown with algorithmically detected spindles masked in blue. Spindles tend to be present within the light off periods. Sigma band (9-16Hz) filtered traces are shown below as well as relative sigma power compared to broadband (1-30Hz) filtered traces above. A spectrogram heatmap is shown of the EEG signal. **f**, 100 randomly sampled 10s segments were drawn from NREM states. Random sampling iterations occurred 100 times. Segments with datapoints greater than 300e^-5^V were excluded from analysis. Average of relative sigma power during VPi (top; *n* = 6 mice 0.5Hz and 1.5Hz, *n* = 7 mice 1Hz) or TRN (bottom; *n* = 6 mice 0.5Hz and 1.5Hz, *n* = 7 mice 1Hz) optostimulation (NREM only) for 0.5, 1, or 1.5Hz. **g**, cross-correlogram for relative sigma power and optostimulation on times. **h**, Characteristics of detected spindles during optostimulation compared to no stimulation for both VPi and TRN optostimulation experiments. n.s. all, Mann-Whitney U test. *P* < 0.05 (*), n.s. = not significant.

We quantified the delay of TC and VPi or TRN peak activity following optostimulation of VPi or TRN neurons. Following VPi activation, TC neuron activity peaked at approximately 0.4759 ± 0.0302s and VPi neurons peaked at 0.3271 ± 0161s. Therefore, TC neurons peaked at approximately 150ms following VPi neurons (*P* = 0.0476; one-sided Mann Whitney U test; **Fig. 4d**). Similarly, TC neuron activity peaked at approximately 0.4351 ± 0.0063s following TRN stim and TRN neurons peaked at 0.2892 ± 0.0109s. TC neurons also peaked at approximately 150ms following TRN neurons (*P* = 0.0286; one-sided Mann-Whitney U test; **Fig. 4d**). These timepoints are in line with the temporal kinetics of T-type calcium channel dynamics and natural bursting tendencies of TC neurons [11] [12].

In individual mice, algorithmically detected sleep spindles were visible immediately after optostimulation of either VPi or TRN neurons (**Fig. 4e**). Locations of detected sleep spindles were also aligned with increases in relative sigma power (proportion of sigma power over the bandpass power; **Fig. 4e**). These trends of relative sigma power seen in individual mice were also apparent in averages across all mice. Peaks of relative sigma power are elicited after each optostimulation of either VPi or TRN neurons at 0.5, 1 or 1.5Hz (**Fig. 4f**) though 1.5Hz may be too fast and is clipping spindles as they occur. 1Hz results summarized in a cross-correlogram show a clear 10 peaks per 10s of lag with VPi activation eliciting a milder correlation as compared to TRN activation (**Fig. 4g**). In contrast, inhibition of VPi failed to induce spindles correlated to optoinhibition whereas TRN optoinhibition led to a delayed peak of relative spindle power (**Extended Data Fig. 7**). We also compared spindle characteristics of algorithmically detected induced spindles versus natural spindles detected in 10 second NREM periods where no optostimulation occurred (**Fig. 4h**). We have found no significant difference in number of spindles per 10s period, spindle duration, amplitude, absolute sigma power, relative sigma power, spindle frequency, number of oscillations, and symmetry between any combinatorial pair of VPi- or TRN-induced spindles versus naturally occurring spindles (**Fig. 4h**).

### VPi neurons regulate spindle dynamics and learning

We have revealed through rabies tracing and optogenetic strategies that TRN neurons provide direct input to both VPi and LGNi neurons (**Extended Data Fig. 8-9**) firmly establishing local thalamic interneurons within the thalamocortical circuit. To investigate the role of this TRN connectivity to VPi neurons and their presumed redundant function in facilitating TC rebound bursting and spindle generation we injected S5E2-ChR2 and DIO-eNpHR into the VP thalamus of Sox14-Cre mice, enabling rhythmic spindle induction through TRN neurons while simultaneously inactivating VPi neurons (**Fig. 5a,b**). On the first recording day, TRN neurons were optostimulated at 1Hz with a light-on of 0.2s during NREM sleep for the first 12 hours. In the subsequent 12 hours, TRN neurons were optostimulated while VPi neurons were also inactivated (**Fig. 5b**). To assess potential circadian influences on spindle properties, a second recording session was conducted for each mouse, reversing the order of stimulation conditions (**Fig. 5b**). Comparison of TRN stimulation alone versus TRN stimulation combined with VPi inhibition revealed a significant increase in spindle duration, relative spindle power (for day 1), and spindle count with VPi inhibition (**Fig. 5c-e**). These results suggest that VPi neurons are key regulators of spindle dynamics by dampening spindle length and occurrence.

**Fig. 5.**
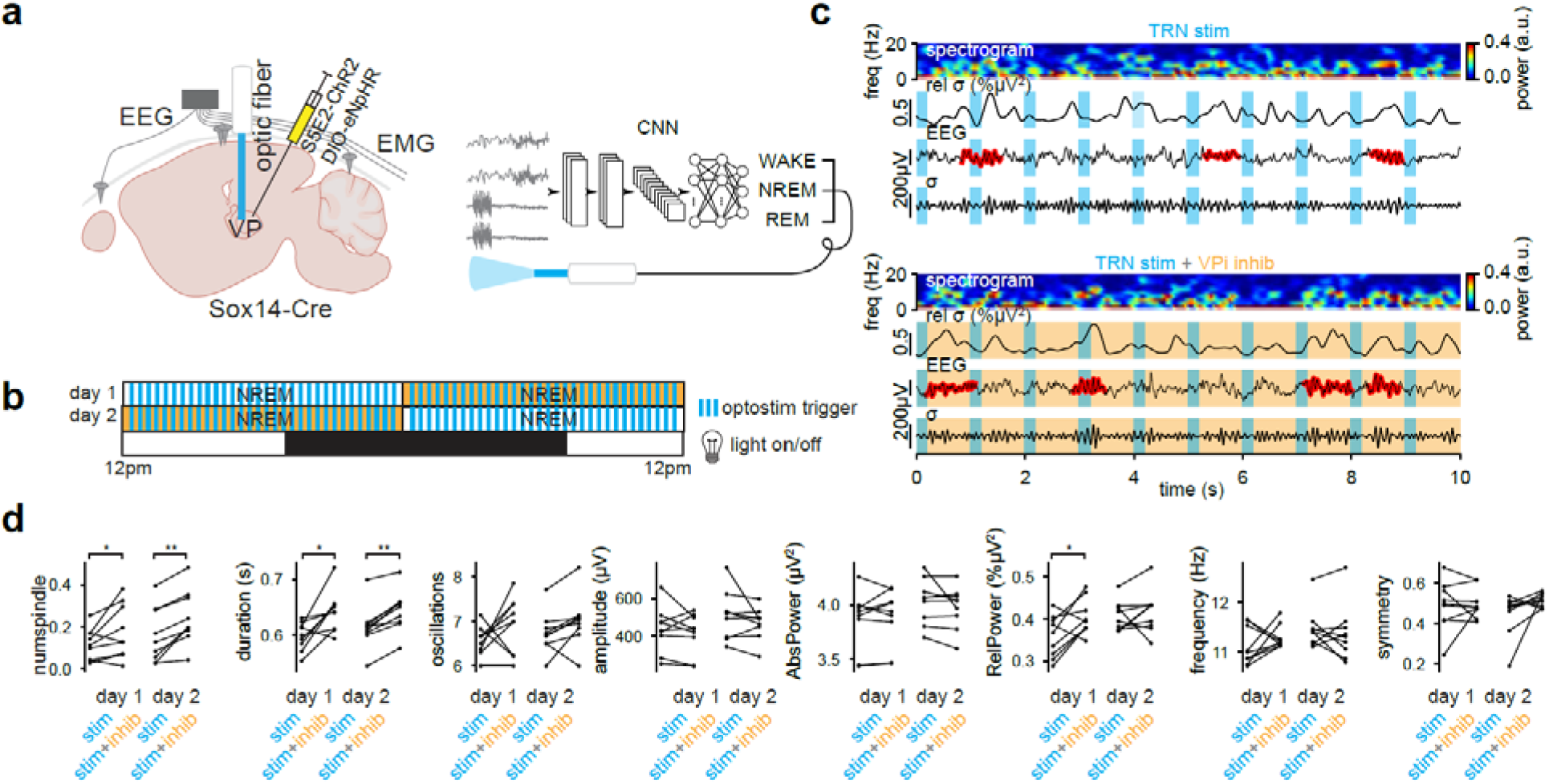
VPi inhibition alters spindle incidence and temporal kinetics. **a**, Stereotaxic injection of S5E2-ChR2 and DIO-eNpHR in Sox14-Cre mice. EEG leads are implanted with screws into left and right somatosensory areas (see Fig. 1). The optic fiber is implanted over the VP. **b**, TRNs are optostimulated in a closed loop design during NREM according to time blocks. 24 hour recordings are split into halves. In the first day of recording, TRN neurons are optostimulated at 1Hz in the first half. In the second half, VPi neurons are inhibited during TRN optostimulation. In the second day of recording, this order is reversed. **c**, Example data from day 1 first and second half experiments. Right somatosensory EEG data is shown with algorithmically detected spindles masked in red. Spindles tend to be present within the light off periods. Sigma band (9-16Hz) filtered traces are shown below as well as relative sigma power compared to broadband (1-30Hz) filtered traces above. A spectrogram heatmap of the EEG signal is shown. **d**, Characteristics of detected spindles for each recording day. The first 70 samples of 10s segments were obtained from NREM states (in which optostimulation occurs). Segments with datapoints greater than 300e^-5^V were excluded from analysis. The same mice were used for day 1 and day 2 (*n* = 9 mice). Day 1: numspindles *P* = 0.0391, duration *P* = 0.0273, oscillations *P* = 0.2626, amplitude *P* = 0.4961, AbsPower *P* = 0.7343, RelPower *P* = 0.0391, frequency *P* = 0.4258, symmetry *P* = 0.8203, Day 2: numspindles *P* = 0.0039, duration *P* = 0.0039, oscillations *P* = 0.0977, amplitude *P* = 0.3594, AbsPower *P* = 0.4961, RelPower *P* = 0.3594, frequency *P* = 0.5703, symmetry *P* = 0.0977, Wilcoxon signed-rank test. *P* < 0.01 (**), *P* < 0.05 (*).

To determine if VPi neurons influence learning through spindle regulation during NREM sleep, mice were then trained on a head-fixed go/no-go whisker detection task, in which they learned to report the presence or absence of a metal rod presented within whisker range (**Fig. 6a**). Correct responses were rewarded with water, while false alarms were penalized with an air puff. During training, EEG and EMG were recorded continuously outside of task periods. Detected NREM sleep epochs triggered continuous LED illumination to inactivate VPi neurons (**Fig. 6b**). All mice performed above chance across training, but differences emerged in the rate of learning between groups (**Fig. 6c**). Control mice displayed a steady improvement in performance across training days, whereas VPi-inhibited mice plateaued at a lower performance. VPi-inhibited mice displayed a significantly lower improvement, calculated as the difference between average performance in the second half versus the first half of training (*P* = 0.0286; **Fig. 6d**). Control mice also tended to reach higher sensitivity (d^′^) and exhibited biases closer to zero, indicating more balanced responses between go and no-go trials (**Fig. 6c,d**). These trends pointed toward reduced detection and greater response bias following VPi inactivation. Together, these results show that VPi inhibition impaired performance improvement and reduced the refinement of decision metrics across training, suggesting that VPi neurons contribute to spindle dynamics during NREM sleep that are necessary for sensory learning.

**Fig. 6.**
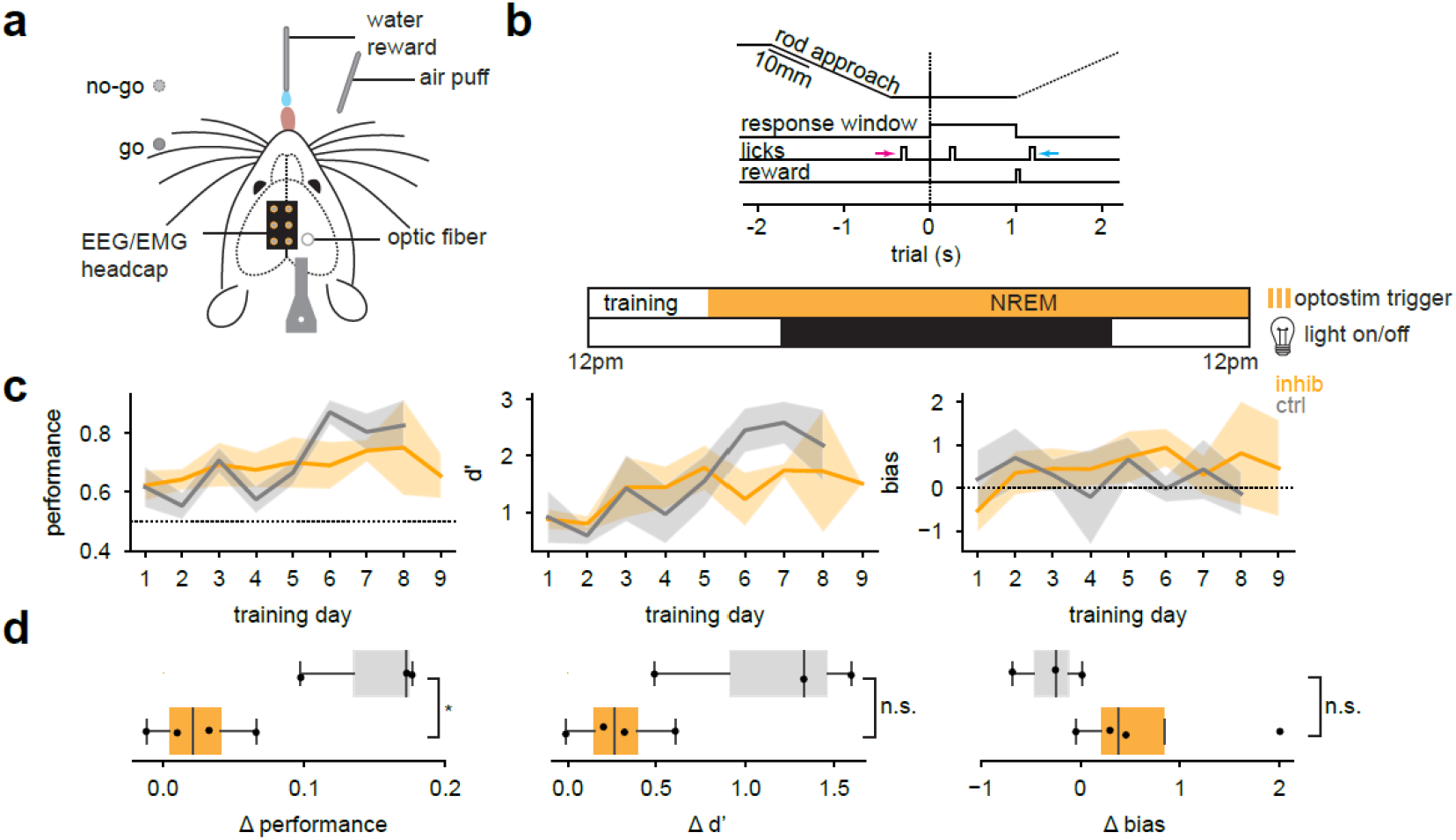
VPi neurons regulate learning through NREM sleep. **a**, Sensory task paradigm. Mice received a go no-go whisker sensory task in which they must detect the presence or absence of a metal rod. Hits were rewarded with water whereas False Alarms trigger an air puff penalty. **b**, The 1 second response window begins 2 seconds into each trial. Early (magenta arrow) and late (cyan arrow) licks were logged but did not trigger rewards or penalties. Penalties are applied immediately, while rewards are given after the reward window. Following daily training, mice are placed in recording enclosures, where detected NREM segments of EEG and EMG trigger solid light. **c**, Performance, d’, and bias metrics for control and optoinhibition groups. The control group includes 2 unperturbed mice and 1 mouse that underwent the same recording and stimulation paradigm as the optoinhibition group. Control mice were pooled, as light alone did not appear to affect learning. Optoinhibition: *n* = 4 mice, control: *n* = 3 mice. **d**, Boxplots of Δperformance, Δd’, and Δbias calculated as the difference in the average values from the second half minus the average values from the first half in which the halfway point is determined as the 4^th^ session from the last training day. Mann Whitney U-test one-sided: Δperformance *P* = 0.0286, Δd’ *P* = 0.0571, Mann Whitney U-test two-sided: Δbias *P* = 0.1143). *P* < 0.05 (*), n.s. = not significant.

## Discussion

In this study, we demonstrated that VPi neurons receive inhibitory input from TRN neurons and actively contribute to thalamocortical circuit dynamics. Brief optostimulation of either TRN or VPi neurons induced spindles entrained to the stimulation frequency. Inhibiting VPi activity during TRN-driven rhythmic stimulation disrupted spindle timing, increasing both their occurrence and duration. Furthermore, inactivating VPi neurons during NREM sleep impaired performance in a whisker-based sensory detection task. Together, these findings indicate that VPi neurons shape the temporal structure of spindle oscillations and are necessary for sleep-dependent sensory learning.

Whole-cell recordings and morphological reconstructions of LGNi and VPi neurons have thus far provided compelling evidence that these neurons share a common cell type [48] [49] [50] [9] [51] [52] [53]. Though the synaptic connection of TRN to LGNi neurons has been a debated topic in previous work [54], we have demonstrated that LGNi and VPi neurons receive inhibition from neighboring TRN neurons. Monosynaptic TRN input supports that both LGNi and VPi neurons are active participants in spindles and perhaps other oscillatory events. The LGN is not often studied in the context of sleep; however, the LGN does generate sleep spindles like other thalamic nuclei [55] [56]. Accordingly, it is likely that the LGNi neurons serve a similar function in the LGN of modulating temporal spindle features as that of the VPi in the VP. Sleep spindles are ubiquitous in the thalamus [57] and the logic of modality-specific localization of spindles is not yet fully understood. It has been shown that spindles facilitate communication between the LGN and the primary visual cortex during sleep, which is essential for consolidating adaptive changes in visual system function following novel visual experiences [58] [59] [60] [61]. Similarly, motor task improvements correlated with an increase in NREM sleep and exposure to a new sequence of motor tasks led to an increase in spindle events [62] [63]. This raises the broader question of how local thalamic interneurons contribute to the tuning of spindle dynamics across sensory modalities. It also invites further exploration into whether the modular architecture of subcortical processing represents an evolutionarily conserved strategy across different sensory systems.

Interestingly, LGNi and VPi share the same embryonic origins within the tectum (both Sox14-positive) and migrate postnatally to reach their final destinations in either thalamic nucleus [48] [49]. Consequently, migrating local interneurons are sensitive to retinal input during development which is essential for their proper localization and integration into the LGN [64]. Similarly, in a transgenic mouse line lacking retinal ganglion cells, the absence of retinal inputs was found to result in rerouting of migrating local interneurons to the VP leading to inappropriate localization and decreased numbers in the LGN [65]. These findings raise the intriguing possibility that local interneurons migrate to regions where they are most needed, adapting to the functional demands of the local circuit. VPi neurons are distributed throughout the VP, forming diverse synaptic connections with TC relay neurons and with each other [9]. This strategic positioning enables them to function as a third component of the thalamo-reticular rhythm generator, uniquely suited to regulate spindle dynamics. It is conceivable that the synaptic connections established during development, define the spatial constraints that govern spindle dynamics later in life. However, the extent of the plasticity of these connections and whether new connections can be formed or existing ones strengthened in adulthood remain open questions.

Through developmentally guided migration, VPi neurons may influence not only the temporal features but also the spatial patterns of sleep spindles, potentially shaping the plasticity rules that govern coordinated learning across somatotopically distinct regions. Given their expansive dendritic morphology, thalamic local interneurons have traditionally been viewed as multiplexors capable of integrating diverse sensory inputs across spatially extensive thalamic areas. However, recent anatomical tracing in LGNi neurons has shown these interneurons are specialized to selectively integrate subsets of retinal inputs [66]. Determining the organization and specificity of somatotopic inputs onto VPi neurons will therefore be essential for clarifying whether local interneurons perform conserved or distinct functional roles in the VP thalamus.

The percentage of local interneurons is known to be larger in the somatosensory thalamus of cats, humans, and other mammals [7] [67]. However, mice have been primarily used as the paradigm for experimentally dissecting fundamental concepts of thalamocortical function. We propose that the relatively low abundance of VPi neurons in mice has obscured their important roles in thalamic circuit physiology. For example, spindle dynamics have traditionally been attributed to the TRN neurons, whose intrinsic properties and connectivity appear to be ideally suited for this role. Our findings demonstrate that this responsibility is, at least in part, shared by VPi neurons. While the precise mechanisms of VPi neuron involvement remain to be fully elucidated, direct electrophysiological recordings from VPi neurons and somatotopic mapping of local interneurons across the VP will be essential for refining our understanding of their role in spindle function. By incorporating thalamic local interneurons into models of spindle dynamics, this work lays the foundation for a more realistic understanding of the thalamocortical circuit and mechanisms of rhythmogenesis.

## Methods

### Animal Models

All procedures were carried out in strict accordance with the US National Institute of Health (NIH) guidance for the care and use of laboratory animals and have been approved by the Animal Care and Use committees at Columbia University. Both males and females were used for this study. The following mouse lines were used: PV-Cre (JAX 017320), C57BL6-J (JAX 000664). The Sox14-Cre mouse line was a kind gift from Dr. Alessio Delogu (King’s College London, UK). Animals were housed on a 12hr light:dark cycle (light on 0700h, light off 1900h) with access to food and water *ad libitum*.

### Ex vivo patch clamp recording

Mice were quickly decapitated under isoflurane sedation. Brains were placed in artificial cerebral spinal fluid (aCSF). Coronal slices (300□µm) containing the ventral posterior (VP) thalamus were cut on a vibratome (Leica VT1200) in sucrose cutting solution containing 2.5□mM KCl, 10□mM MgCl_2_, 0.5□mM CaCl_2_, 1.25□mM NaH_2_PO_4_, 26□mM NaHCO_3_, 234□mM sucrose, 11□mM glucose. Slices were then transferred to aCSF containing 126□mM NaCl, 26□mM NaHCO_3_, 2.5□mM KCl, 1.25□mM NaH_2_PO_4_, 2□mM CaCl_2_, 1□mM MgCl_2_, and 10□mM D-glucose bubbled with 95% O_2_ and 5% CO_2_. Slices were incubated at 34□±□1□°C for 30□min before resting at room temperature prior to the experiment. Regular pipette intracellular solution contained 127□mM potassium gluconate, 8□mM NaCl, 4□mM ATP-Mg, 0.6□mM EGTA, 0.3□mM GTP, 10 HEPES, and 8.1□mM biocytin adjusted to pH 7.3–7.4 with KOH. Osmolarity was adjusted to 290–300□mOsm. Local interneurons of the VP or LGN were located using the presence of mCherry or GFP fluorescence. Cells were targeted approximately 20-100µm from the slice surface. Patch-clamp Luigs and Neumann micromanipulators and stage were controlled using the Remote Control SM-10 Touch (Luigs and Neumann). Recordings were performed in whole-cell mode with MultiClamp 700B amplifiers (Molecular Devices) in the current clamp mode at 34 ± 1°C bath temperature.

Data acquisition was performed through an Axon Digidata 1550B (Molecular Devices) connected to a PC running pClamp 11 (Molecular Devices). Recordings were sampled at 10KHz and filtered with a 2-kHz Bessel filter. Patch pipettes were pulled with a Flaming/Brown micropipette puller P-80PC (Sutter Instruments) and had an initial resistance of 3-8MΩ. Series resistance was automatically compensated and was typically around 15MΩ or lower. The membrane potential values given were not corrected for the liquid junction potential which was approximately - 16mV.

### Interface recording

Slices were placed in an interface recording chamber and maintained at 34 ± 1 °C bath temperature. Neuronal activity was recorded with commercially available linear array probes (Model: A16x1-2mm-100-177, NeuroNexus, Ann Arbor, MI, USA) with 16 contacts (spacing: 100µm distance), spanning 1500µm. Probes were slowly lowered across the VP thalamus. An optic fiber delivering 595nm or 465nm light was placed directly above the VP. Data acquisition was performed through a digitizing board (SI-4, Tucker-Davis Technologies) connected to a real-time acquisition processor (RZ10x, Tucker-Davis Technologies) and PC workstation (WS-8, Tucker-Davis Technologies) running custom-written routines in Synapse (Tucker-Davis Technologies). Optogenetic light sources were integrated in the RZ10x and controlled through the Synapse software. Recordings were sampled at 24 kHz. Spike sorting was performed using the integrated PCA (principal component analysis) spike sorting tool within Synapse. Candidate spikes were detected using a threshold set at 5 standard deviations above the root mean square (RMS) of the baseline signal. During an initial training period, candidate spike waveforms were collected and used to compute the first three principal components capturing the greatest variance for each recording channel. Subsequent spike waveforms were projected into this three-dimensional principal component feature space, forming distinct clusters. These waveform clusters were then classified using a Bayesian clustering algorithm, isolating waveforms originating from individual neurons.

### Ex vivo 2-photon imaging

2-photon images were acquired on an acousto-optic deflector (AOD)-based microscope (Femto 3D Atlas, Femtonics) using a 20x water-immersion objective (Olympus XLUMPLFLN, 1.00 NA, 2.0 mm WD). Slices were placed in a heated recording chamber (Luigs and Neumann) perfused with oxygenated artificial cerebrospinal fluid (aCSF) at 34 ± 1 °C. A concentric bipolar stimulating electrode (WPI) was positioned in the VP thalamus ∼100 μm from the recorded region, and stimulation pulses (300 μA, 0.2 ms, 0.5-20Hz) were delivered via an isolated stimulator (A-M Systems). GCaMP fluorescence was excited with brief pulses of a femtosecond laser (Chameleon, Coherent) tuned to 940 nm at ∼10–40 mW power at the sample plane. Images were acquired using MES software (Femtonics). Data were analyzed using custom Python code, with an analysis pipeline including background subtraction, photobleach correction by fitting and dividing a single-exponential function to mean fluorescence, running mean filtering (5-point window), and Δf/f calculation using a pre-stimulus baseline.

### Immunohistochemistry

Mouse brains were perfused with 4% paraformaldehyde for sectioning. Brains were cryosectioned at 30µm. Sections were stained with a Gad67 antibody (1:2000, Sigma) and tagged with a secondary antibody reporter (goat anti-mouse 1:500, Invitrogen). Sections were then mounted onto slides and imaged with a Zeiss 810 confocal microscope. Colocalization analysis was performed manually using ImageJ. Identification of brain regions for analysis of monosynaptic tracing data was performed by overlaying the Allen Institute Adult Mouse Brain Reference Atlas [68] to each slice.

### Stereotaxic surgery

Mice were anesthetized using a mixture of ketamine and xylazine (100mg/kg, 10mg/kg, intraperitoneally) and placed on a stereotaxic base with a heating pad. Following asepsis, an incision is made on the scalp to expose the skull. Bregma is located and craniotomies are performed above the area of interest (VP thalamus, AP: -1.5mm, ML: ±1.8mm, DV: -3.5mm). 400-600nL of a solution containing viral constructs are injected into the region of interest using a nanosyringe (WPI, Nanofil 10µL) controlled by a microinjector (WPI, Micro2T SMARTouch). To perform monosynaptic rabies tracing, we used a Cre-dependent helper virus, AAV1-hSyn-FLEX-TVA-P2A-eGFP-2A-oG (Salk Institute), to selectively express the TVA receptor and rabies glycoprotein (oG) in Cre-expressing neurons. Following injection, the AAV virus was allowed to incubate for two weeks. EnvA G-deleted Rabies-mCherry (Salk Institute) was injected into the same location at an injection volume of 100-200nL to infect TVA-expressing neurons and allow retrograde transsynaptic labeling. Mice were perfused 7 days post-rabies injection.

If mice were to be recoded for *in vivo* fiber photometry or optogenetics studies, mice were implanted with an optic fiber and EEG/EMG leads one week post viral infection. An optic fiber (diameter 400µm, 0.66NA, Doric) was lowered to approximately 100µm above the VP thalamus. For EEG implantations, a ground screw was placed near the olfactory bulb and a reference screw in the cerebellum. Recording screws were also placed in the left and right somatosensory cortex (AP: -1mm, ML: ± 2mm). Two EMG electrodes were placed into the trapezius neck muscle. All implants were secured into place with dental cement (Metabond). All EEG and EMG wires were connected to a 6 pin millmax piece. Animals were returned to their home cage and allowed to recover for at least a week prior to any further experimentation.

### Linear Classifier

For each mouse, simultaneous fiber photometry (GCaMP) and electrophysiological recordings were segmented into consecutive 10s epochs. Data were analyzed separately for two classification tasks: (1) predicting movement state from EMG and GCaMP during wakefulness, and (2) predicting spindle occurrence from EEG and GCaMP during NREM sleep. GCaMP signals were downsampled to 50 Hz with low-pass filtering at 1 Hz. For both classifiers, we extracted a consistent feature set from the downsampled GCaMP traces: standard deviation (std), median absolute deviation (mad), interquartile range (iqr), root mean square of successive differences (rms_dx), and relative power in the 0.2–0.5 Hz frequency band normalized by the total spectral power between 0.05–3 Hz (pow_0.2_0.5). For the movement classifier, ground truth labels were derived from the EMG signal. The rectified EMG was smoothed to generate an envelope, and movement epochs were defined using a threshold equal to the median plus seven times the robust standard deviation (mad). 10s epochs exceeding this threshold were labeled as “movement,” while all other epochs were labeled as “no movement.” For the spindle classifier, labels were assigned based on automatic detection of spindles in the EEG, using the spindle detection algorithm described below. Each 10s epoch was thus classified as either “spindle” or “no spindle.” Classification models were implemented in Python using scikit-learn [69]. Models were constructed as a pipeline that included feature standardization followed by logistic regression. A logistic regression with elastic net regularization was chosen to balance sparsity (L1 penalty) and stability (L2 penalty). To evaluate generalization across animals, leave-one-mouse-out (LOMO) cross-validation was employed using scikit-learn’s LeaveOneGroupOut() function. On each fold, data from all but one mouse were used for training, and the withheld mouse was used for testing. Model performance was quantified by the receiver operating characteristic (ROC) area under the curve (AUC). To establish baseline chance levels, labels were randomly shuffled within each mouse, and classifiers re-evaluated under the same cross-validation framework.

### Convolutional Neural Network Training for Sleep State Classification

Approximately 1800hrs of EEG and EMG signals were preprocessed and used to train a deep learning model for automatic sleep stage classification. Downsampled signals containing two channels of EEG (right and left somatosensory cortex) and two channels of EMG (right and left trapezius muscle) were loaded from EDF files using the MNE-Python library [70], followed by Z-score normalization using Scikit-Learn to standardize the data. The normalization scalers were saved and later applied during inference to ensure consistency between training and evaluation. Data were segmented into epochs of 10s windows. We implemented a 1D convolutional neural network (CNN) using PyTorch, trained on this multichannel time-series data. Prior to model training, the dataset was partitioned into training, validation, and test sets (70-15-15%) and mini-batch data loaders were created with a batch size of 256 without shuffling to maintain temporal structure. Feature extraction was performed through two 1D convolutional layers each followed by ReLU activation and dropout (p=0.5) to reduce overfitting. The extracted features were flattened and passed through a fully connected layer followed by a final classification layer mapping features to three sleep stages (wake, NREM and REM). To address class imbalance due to the relative scarcity of REM events, class weights were computed based on the inverse frequency of each class in the training set and incorporated into the cross-entropy loss function. The model was trained using mini-batch gradient descent for up to 100 epochs, with the Adam optimizer applied to update model parameters. At each epoch, training and validation losses were recorded, and model performance was evaluated using overall accuracy and class-wise accuracy, computed from a confusion matrix. The entire training process was executed on a GPU (CUDA-enabled), ensuring computational efficiency and optimized performance. After training to approximately a 95% prediction accuracy, the model was used for automated sleep classification of unseen EEG/EMG recordings in real-time or offline sleep scoring.

### Fiber Photometry

To habituate mice to the recording conditions, mice were habituated to the enclosure for one day, followed by habituation with recording cables (fiber optic and EEG/EMG cables) for another day. Daily monitoring and experiment set-up occurred at approximately noon. Calcium dependent GCaMP fluorescence was measured using a sinusoidal modulated LED (465nm 330Hz, 405nm 210Hz, Tucker Davis Technologies) and detected using an integrated photosensor photodiode (RZ10x, Tucker Davis Technologies). EEG and EMG signals were acquired through a digitizing board (SI-4, Tucker-Davis Technologies) connected to a real-time acquisition processor (RZ10x, Tucker-Davis Technologies) and PC workstation (WS-8, Tucker-Davis Technologies) running custom-written routines in Synapse (Tucker-Davis Technologies). The Δf/f was calculated as: (f-f0)/f0, where f0 was the fitted 405□nm signals. For closed-loop optogenetic stimulation with simultaneous fiber photometry recordings, a 10s rolling buffer of photometry data, acquired via a P08e card (Tucker-Davis Technologies), was used to trigger optogenetic stimulation. Real-time behavioral state classification was performed using predictions from our neural network, which triggered stimulation via the LED driver (Doric) based on classified states. The trained model was loaded from a pre-saved checkpoint, set to evaluation mode, and executed on a GPU (CUDA-enabled). Raw electrophysiological recordings were downsampled and normalized using pre-trained scalers from the training phase to ensure consistency across datasets. Light stimulation patterns were generated in Synapse, and all data analysis was conducted using custom Python scripts.

### Behavioral Apparatus

The sensory task training was conducted within a custom-built behavioral apparatus. The exterior was comprised of a custom aluminum scaffolding housing a blackout curtain and an aluminum breadboard base (Thorlabs). A stepper motor (Pololu 1204) that rotated a 5mm outer diameter metal rod was fitted onto a 30mm linear actuator (Actuonix L12-30-50-6-R) that brought the metal rod to the mouse’s whiskers. Water rewards (5-10µL) were delivered via pump through a blunted stainless steel tube (Luigs and Neumann). Licks were detected through capacitive touch sensors (MPR121, Sparkfun) and penalties were delivered through a solenoid valve (Amazon, B0BWDR2L7S) connected to an oxygen tank. At the start of each trial, house lights were illuminated such that the mouse never fully dark adapts (Amazon, B0C7816B6W). Mice were allowed to run while head fixed on a treadmill to reduce stress (Luigs and Neumann). All components were controlled through an Arduino Uno connected to a dedicated PC via usb. Trial structures were written within the Arduino IDE and trial data was collected in real time using custom python code.

### Go No-Go Training

Mice were water deprived and received their daily water requirements through the sensory task. Mice were closely monitored and weighed to ensure sufficient hydration. *Ad libitum* water was provided if necessary.

Mice learned to detect the presence or absence of a metal rod through a go no-go design. The trials were delivered at random to the mice in equal numbers. Mice progressed through the training in three steps: first the mice were given the randomized go no-go task with automated rewards for every go trial, then the mice underwent the same design with the addition of an air puff penalty for false alarms, and in the final stage, mice were given the randomized task with rewards only for correct responses and air puff penalties for errors. Sessions were terminated when mice became fully satiated, defined as a failure to lick during 10 consecutive go trials, with no licks during the intervening no-go trials (called the disengagement index). It was sometimes necessary to revert mice to previous stages of training, but importantly, only the final stage of training was used in the analysis. In most cases, mice were able to become fully trained through all stages within 14 days with ∼7-9 days in the final stage. Performance was calculated as the number of hits plus correct rejections over all go and no-go trials. d’ was calculated as z-score corresponding to the hits minus the z-scores of the false alarms..

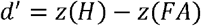

Bias was calculated as the z-score corresponding to the hits plus the z-score corresponding to the false alarms divided by 2

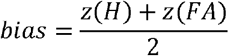

### Sleep Spindle Detection

Sleep spindles were detected using the YASA Python package [71]. The following describes the detection algorithm within this package. First, the raw signal was preprocessed by applying a broadband frequency filter (1–30 Hz). Spindle detection employed three distinct thresholds based on relative sigma power, Pearson’s correlation coefficient, and root mean square (RMS) amplitude, along with duration constraints. Relative sigma power (9–16 Hz) was calculated as the ratio of sigma power to total broadband power using a short-time Fourier transform (STFT) computed over consecutive 2s windows with a 0.2s step size. Relative sigma power calculations in this work outside of sleep spindle detection used a 0.5s window with a 0.1s step size. A relative sigma power threshold was set at 0.2. Next, a correlation threshold was implemented by filtering the raw signal into the sigma frequency range and calculating Pearson’s correlation coefficient between the broadband and sigma-filtered signals within moving windows of 0.3s, with a step size of 0.1s. The correlation coefficient threshold was set at 0.65. Finally, RMS amplitude was calculated using a moving window of 0.3s with a step size of 0.1s. The RMS threshold for spindle detection was defined as the mean RMS plus 1.5 times the standard deviation of the RMS. A soft threshold is calculated by smoothing the decision vector with a 0.1s window. Subsequently, spindle start and end points are defined by identifying indices in the decision vector where at least two out of the three thresholds are exceeded. Spindles that were spaced less than 0.1s apart were merged and spindles that were too short (<0.5s) or too long (>2s) were removed. The same thresholds and constraints were applied for the detection of mu events except for the frequency range which was 8-12Hz.

The final step of the spindle detection algorithm involves extracting key characteristics of each detected sleep spindle. The calculation of spindle start and end times, and consequently duration, is briefly described above. RMS (µV) amplitude is computed across the entire spindle duration. Median absolute power (log10 µV^2^) is determined as the median of the logarithmic sigma-band power derived from the Hilbert transform of the sigma-filtered EEG. Median relative power (% µV^2^) is computed as the median of the relative sigma-band power ratio calculated at each time point within the spindle. Spindle amplitude (µV) is quantified as the peak-to-peak amplitude (maximum minus minimum) using NumPy’s ptp function. Median frequency (Hz) is extracted as the median of instantaneous frequency values, calculated using the Hilbert transform of the sigma-filtered EEG. The number of oscillations within each spindle is quantified by counting peaks in the spindle waveform using SciPy’s find_peaks function, enforcing a minimum interval of 60ms between peaks. Spindle symmetry is represented on a normalized scale from 0 to 1, where 0 indicates the spindle’s start and 1 its end, reflecting the temporal position of the most prominent peak within the spindle.

## Data Visualization

Boxplots were generated to visualize the distribution of data, highlight potential outliers, and overlay individual data points. Data were plotted using Python’s Matplotlib and Seaborn libraries. Each boxplot displayed the median, interquartile range (IQR), and whiskers extending to 1.5 times the IQR. Outliers, defined as values beyond this range, were plotted individually as separate points. To enhance data transparency, all individual data points were overlaid using a jittered scatter plot to reduce overlap. Bar plots were used to display group means, with error bars representing the standard error of the mean (SEM). Individual data points were overlaid where appropriate, also with a jitter, to illustrate the distribution of values within each group. Individual data points represented mice, except for Fig. 1i (Right) and Fig. 2f (Left) in which data points represented averaged folds of the leave-one-mouse-out cross-validation and Fig. 3 in which data points represented sorted neurons.

## Statistical Analysis

All statistical analyses were performed using non-parametric tests to account for potential deviations from normality in the data distribution. For comparisons between two independent groups, the Mann-Whitney U test was used, while for paired data, the Wilcoxon signed-rank test was applied. When comparing more than two groups, the Kruskal-Wallis test was used, followed by post hoc pairwise comparisons with Dunn’s test. The statistical significance threshold was set at *P* < 0.05. All statistical tests were performed using Python (SciPy).

## Acknowledgements

We are grateful to Dr. Alessio Delogu (King’s College London, UK) for the generous gift of the Sox14-Cre mouse line. We would like to thank Dr. Damian Williams and Dr. Franck Polleux for their help in editing this manuscript. J.S. was supported by the NIH BRAIN Initiative grant F32H130023 and the Swiss National Science Foundation Early PostDoc.Mobility grant. T.K.R was supported by the NSF GRFP DGE-2036197. C.D.M was supported by the NIH NINDS grant R00NS104215.

## Author Contributions

Conceptualization, J.S., C.D.M.; methodology, J.S.; Investigation, J.S., T.K.R., C.M., S.L., E.A.S.; writing— original draft, J.S., C.D.M., writing—review & editing, J.S., T.K.R., C.M., S.L., E.A.S, C.D.M..; funding acquisition, J.S., T.K.R, and C.D.M.; resources, C.D.M.; supervision, J.S. and C.D.M.

## Additional Information

**Extended Data Fig. 1–8**

