## Supplemental figures for "Local thalamic interneurons drive spindle termination and enable sleep-dependent learning"

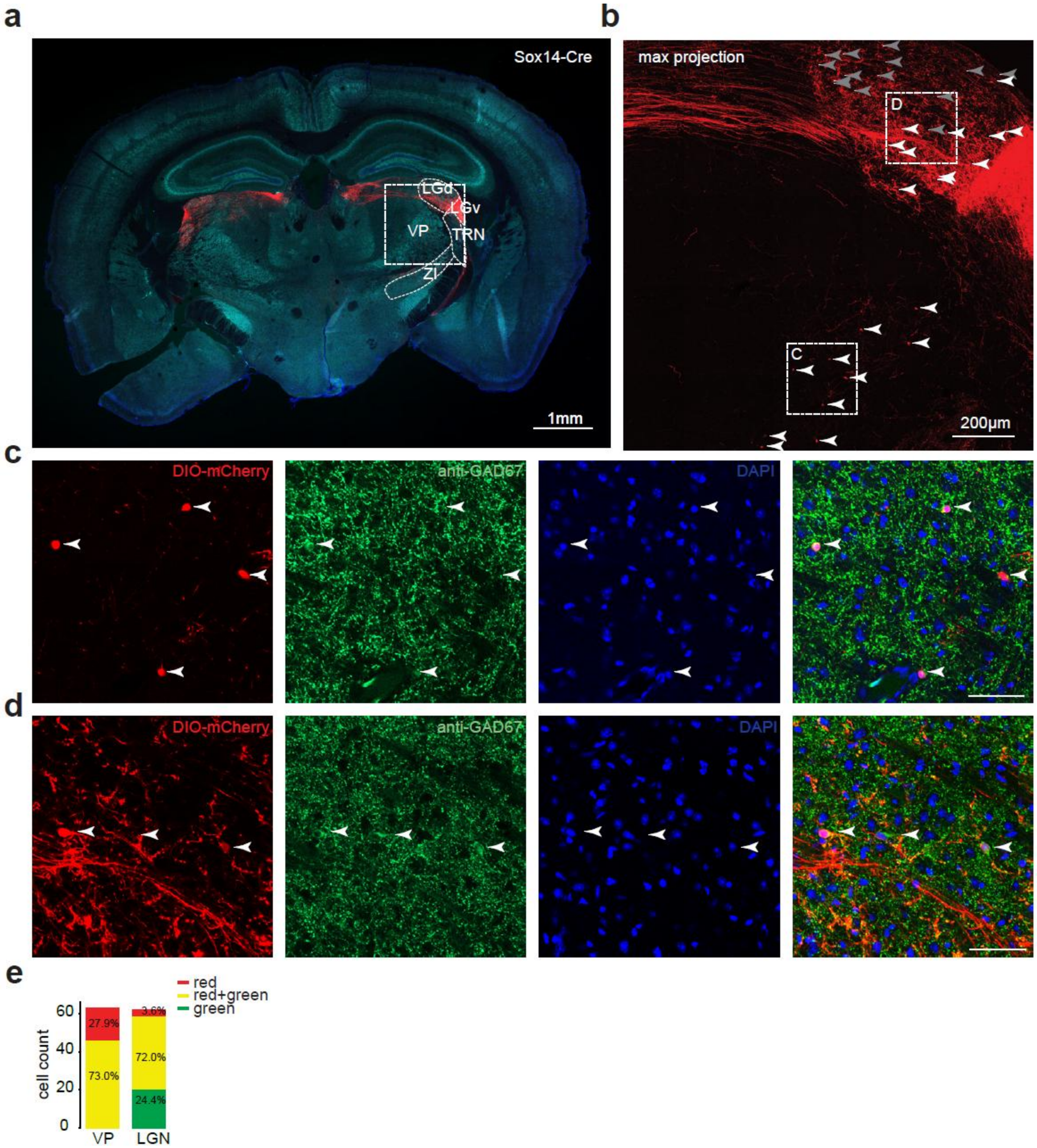


**Extended Data Fig. 1|VPi and LGNi neurons express GAD67.**

**a,** 10x magnification tiled confocal image of mouse brain slice with anti-GAD67, DAPI and virally infected Cre-dependent mCherry in a Sox14-Cre mouse. **b,** Z-stack max projection zoom of inset shown in left panel. Arrows point to detected VPi somas and LGNi somas. Grey arrow point to somas detected with anti-Gad67 only. **c,** Zoom of inset shown in b with channels separated. Arrows pointed to soma of VPis. Combined channels in final panel. **d,** Same as C for somas detected in the LGN. LGNi in the middle is green only. **e,** Proportion of cells that express mCherry only (red) or mCherry and anti-GAD67 (red + green) or anti-Gad67 only (green) for the VP and LGN. VP *n* = 63 cells, LGN *n* = 82, *n* = 2 mice. LGd: dorsal lateral geniculate nucleus, LGv: ventral lateral geniculate nucleus, ZI: zona incerta.

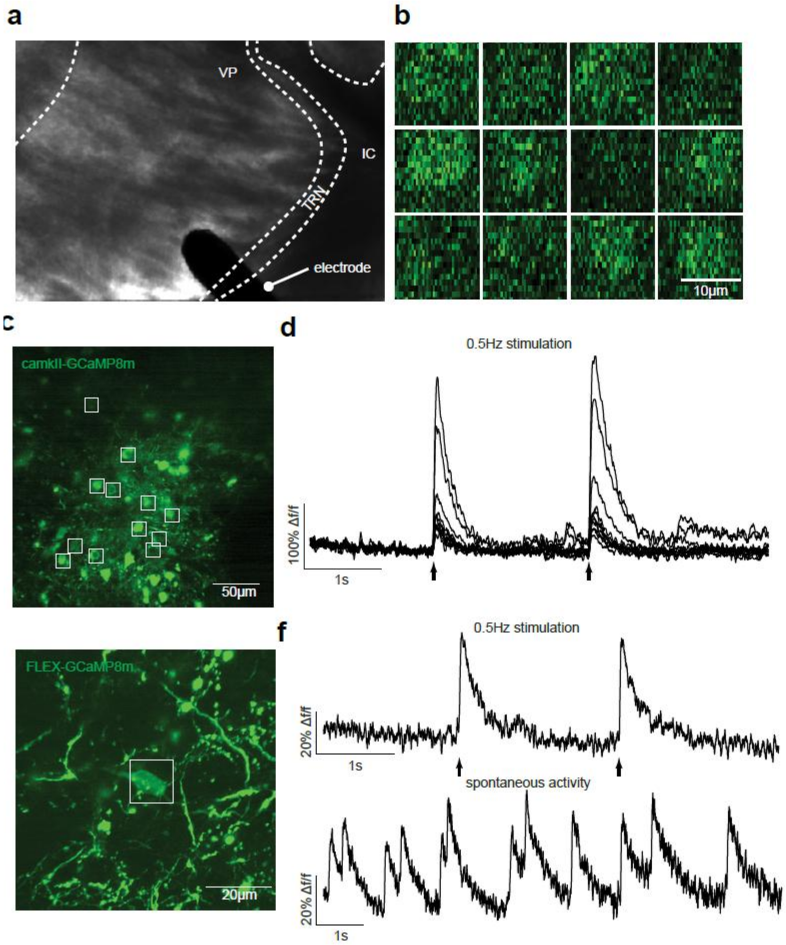


**Extended Data Fig. 2|TC and VPi neurons express GCaMP8m.**

**a,** Slice image of panels b-f. **b,** Chessboard scans of 12 TC ROIs. **c,** Image of GCaMP8m expressed in TC neurons. **d,** GCaMP responses of C. Bipolar electrical stimulation shown with arrows. **e,** Image of GCaMP8m expressed in VPi neuron. **f,** GCaMP response from electrical stimulation (arrows) and subsequent spontaneous activity. AAV9-CaMKII-GCaMP8m: *n* = 1 mouse, AAV9-FLEX-GCaMP8m: *n* = 1 mouse.

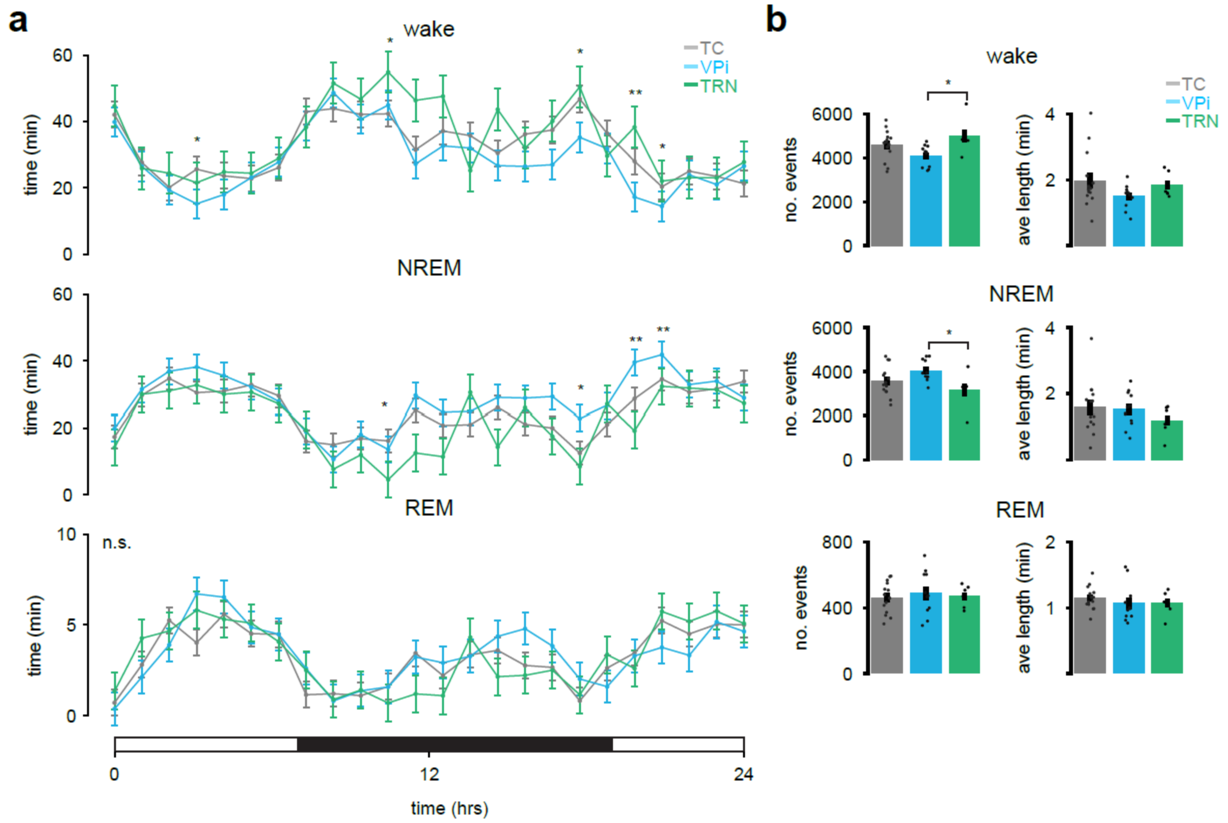


**Extended Data Fig. 3|Sleep characteristics of TC, VPi, and TRN targeted mice.**

**a,** Time spent in wake, NREM, and REM sleep over a 24-hour period for TC, VPi, and TRN-targeted groups. **b,** Number of events and average event length across sleep states for TC, VPi, and TRN-targeted groups. Data represent mean ± SEM. Statistical comparisons were performed using the Kruskal-Wallis test followed by Dunn's multiple comparison test. $P < 0.05$ (*), $P < 0.01$ (**). n.s. = not significant. TC $n = 14$ mice, VPi $n = 11$ mice, TRN $n = 7$ mice.

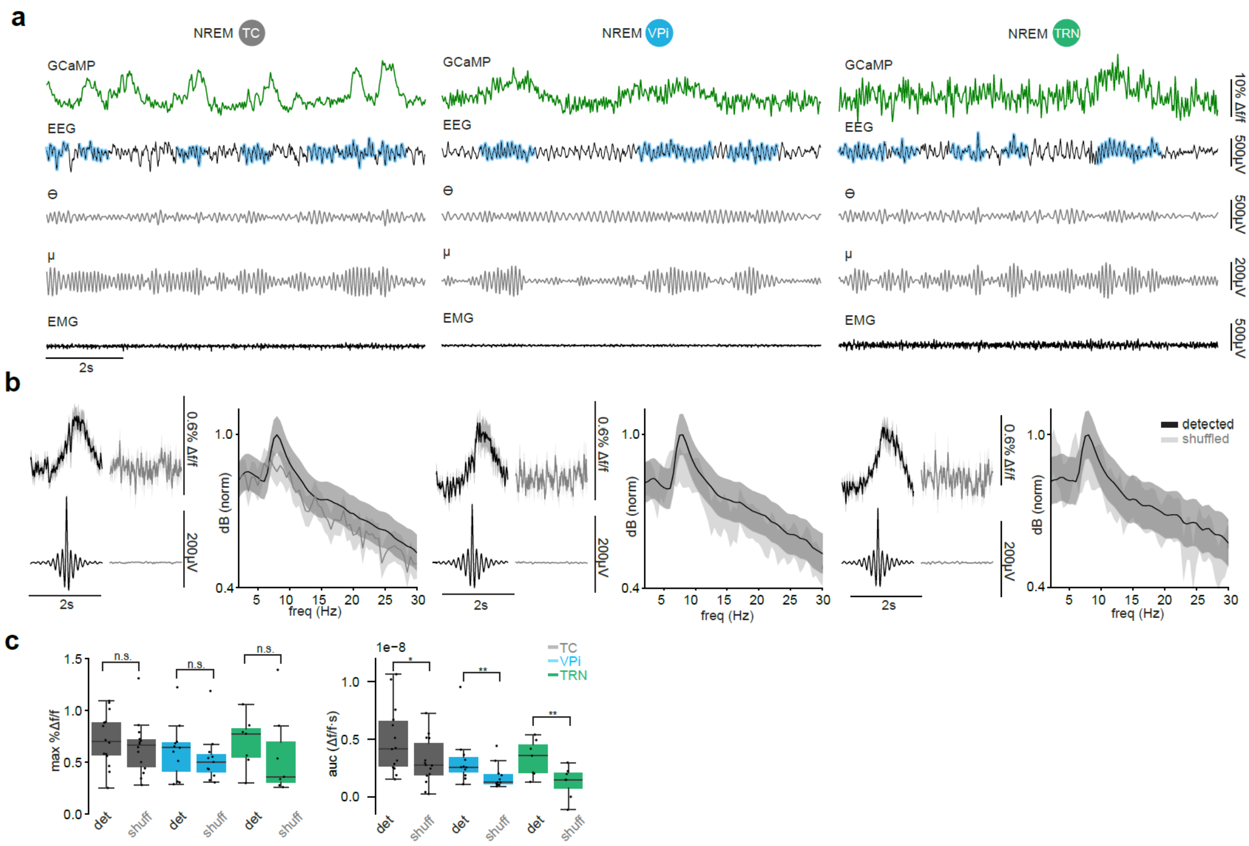


**Extended Data Fig. 4|TC VPi and TRN neurons participate in mu events.**

**a,** Example GCaMP signal, right somatosensory EEG, theta band filtered EEG, mu band filtered EEG, and EMG. Algorithmically detected mu events shown with a blue mask. **b,** Comparing 100 randomly sampled 2s averaged GCaMP activity and EEG detected mu events vs. 100 shuffled 2s segments of REM recordings. Random sampling iterations occurred 100 times. EEG or GCaMP segments with mean values exceeding a z-score threshold of 5 were excluded from analysis. Power spectral density plots of detected vs. shuffled 2s segments of EEG data. **c,** Comparison of calcium responses between detected ("det") and shuffled ("shuff") events across TC, VPi, and TRN populations. Left: Maximum %Δf/f for detected and shuffled events. Right: Area under the curve (auc) of Δf/f responses for detected and shuffled events. Box plots display the median, interquartile range, and individual data points. Statistical comparisons were performed using the Wilcoxon signed-rank test (one-sided) to assess whether detected events were significantly greater than shuffled controls. $P < 0.001$ (***), $P < 0.01$(**), $P < 0.05$(*), n.s. = not significant. TC $n$ = 14 mice, VPi $n$ = 11 mice, TRN $n$ = 7 mice. Data represented as mean ± SEM.

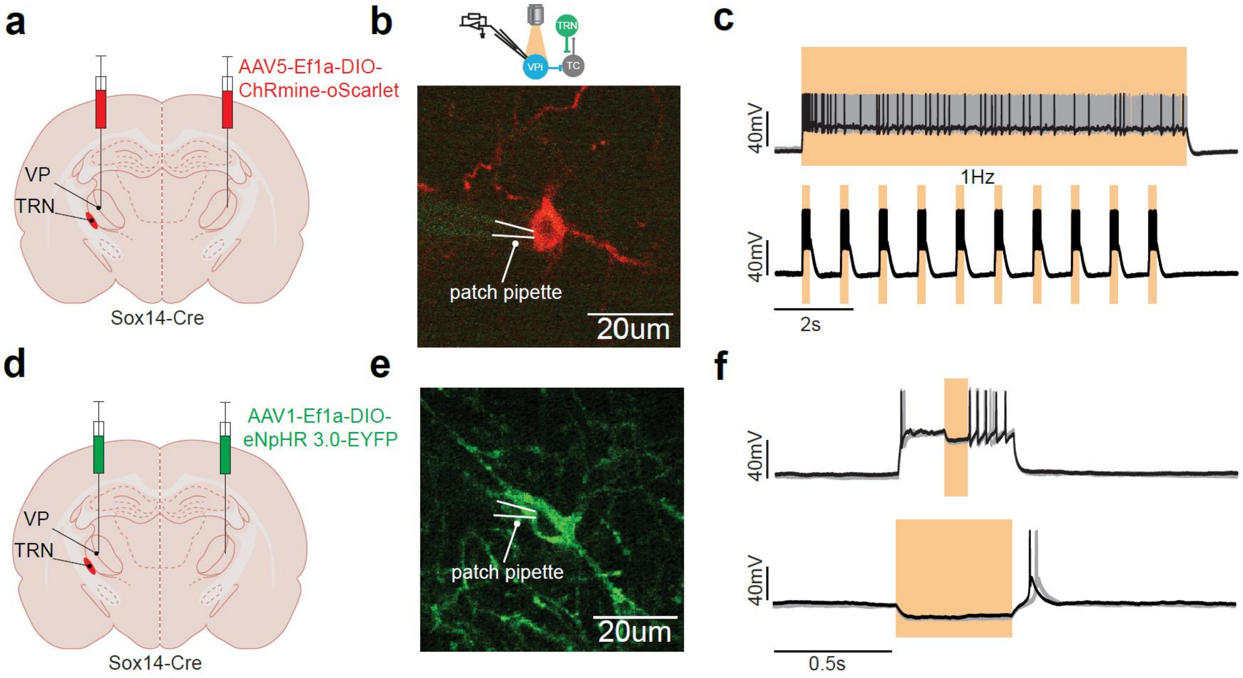


**Extended Data Fig. 5|VPi neurons express ChRmine and eNphR.**

**a,** Stereotaxic injection of AAV5-Ef1a-DIO-ChRmine-oScarlet (bilateral) into the VP of Sox14-Cre mice. **b,** 2 photon image of whole cell patched VPi. **c,** Example of VPi activity during various light protocols. **d,** Stereotaxic injection of AAV1-Ef1a-DIO- eNpHR 3.0-EYFP (bilateral) into the VP of Sox14-Cre mice. **e,** 2 photon image of whole cell patched VPi. **f,** Example of VPi inhibition during various light protocols. ChRmine $n$ = 3 neurons, eNphR $n$ = 6 neurons.

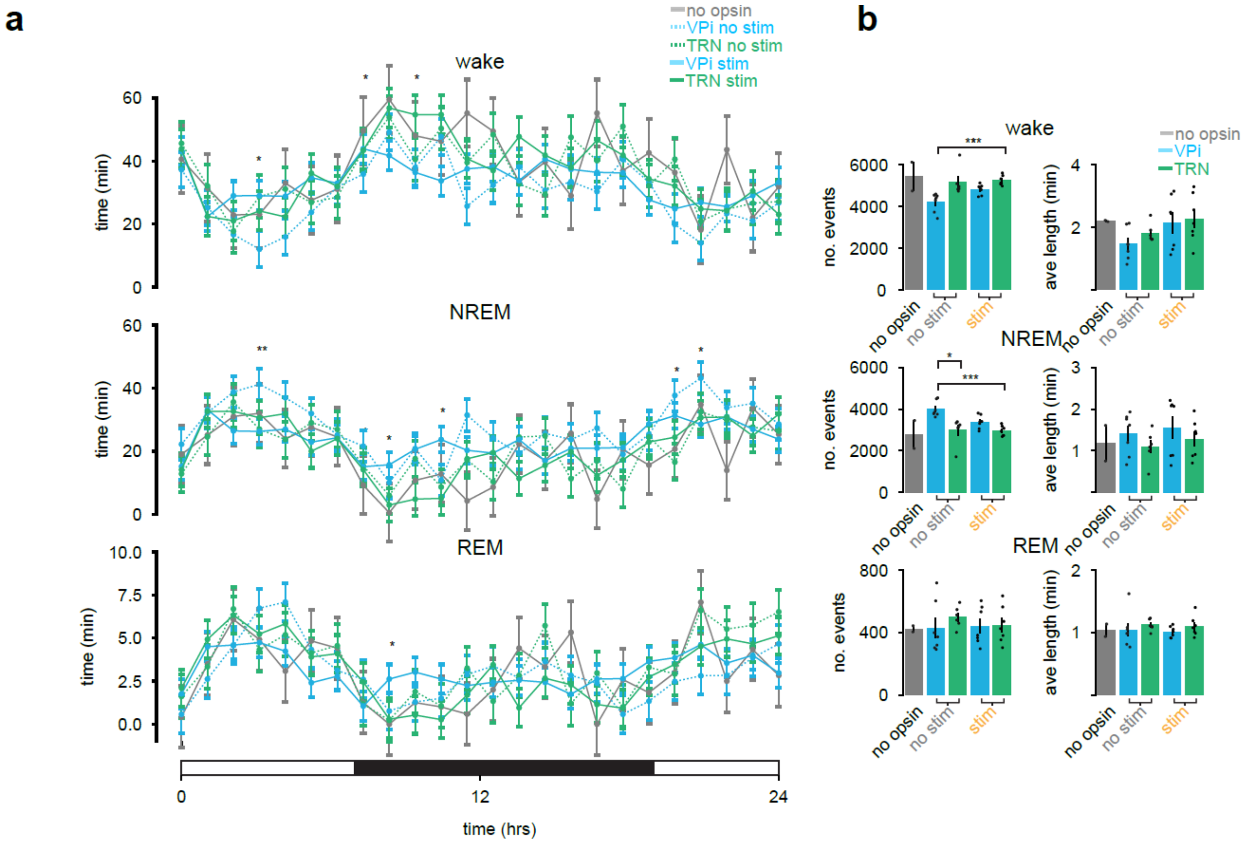


**Extended Data Fig. 6|Sleep characteristics of optogenetically stimulated mice.**

**a,** Time spent in wake, NREM, and REM sleep over a 24-hour period for no opsin control, DIO-chRmine-injected Sox14-Cre mice baseline mice (VPi no stim), DIO-chRmine-injected Sox14-Cre mice with optogenetic stimulation (VPi stim), DIO-chRmine-injected PV-Cre baseline mice (TRN no stim), and DIO-chRmine-injected PV-Cre mice with optogenetic stimulation (TRN stim) groups. **b,** Number of events and average event length across sleep states for mouse groups. Data represent mean ± SEM. Statistical comparisons were performed using the Kruskal-Wallis test followed by Dunn's multiple comparison test. $P < 0.05$ (*), $P < 0.01$ (**), $P < 0.001$ (***), n.s. = not significant. no opsin $n = 2$ mice, VPi no stim $n = 7$ mice, VPi stim $n = 7$ mice, TRN no stim $n = 6$ mice, TRN stim $n = 7$ mice.

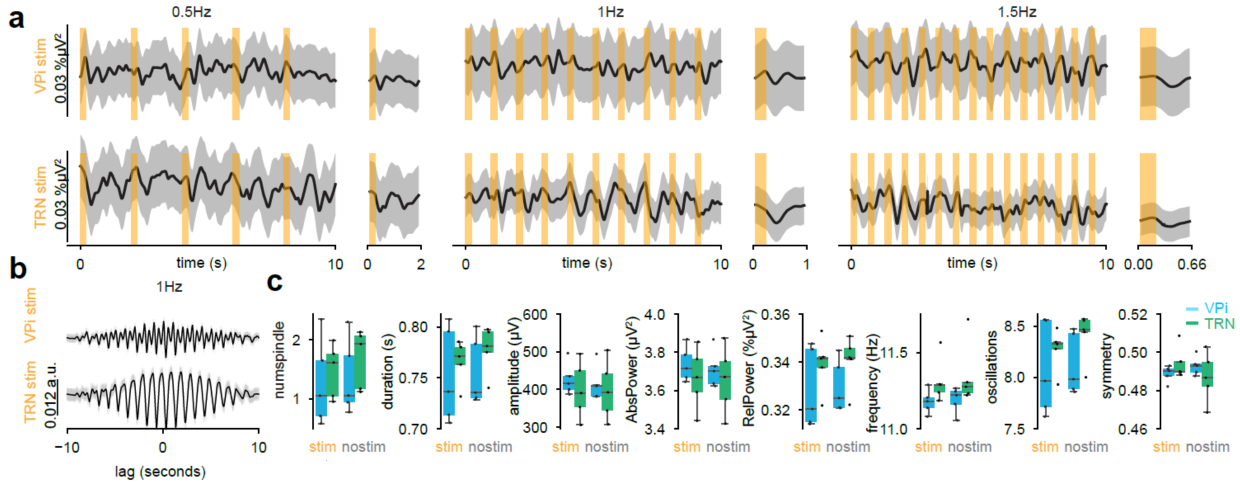


**Extended Data Fig. 7|VPi and TRN inhibition is less effective in entraining sleep spindles.**

**a,** Sox14-Cre or PV-Cre mice have been injected with AAV1-Ef1a-DIO-eNpHR-eYFP. VPi or TRN neurons were optoinhibited according to the same protocool as in Figure 5. 100 randomly sampled 10s segments were drawn from NREM states. Random sampling iteratures occurreed 100 times. Average of relative sigma power during VPi (top: *n* = 4 mice 0.5Hz and 1.5Hz, *n* = 5 mice1Hz) or TRN (bottom: *n* = 4 mice 0.5Hz and 1.5Hz, *n* = 5 mice 1Hz) optoinhibition (NREM only) for 0.5, 1, or 1.5Hz. **b,** cross-correlogram for relative sigma power and optoinhibition on times. **c,** Characteristics of detected spindles during optoinhibition compared to no inhibition for both VPi and TRN optoinhibition experiments. n.s. all, Mann-Whitney U test. n.s. = not significant.

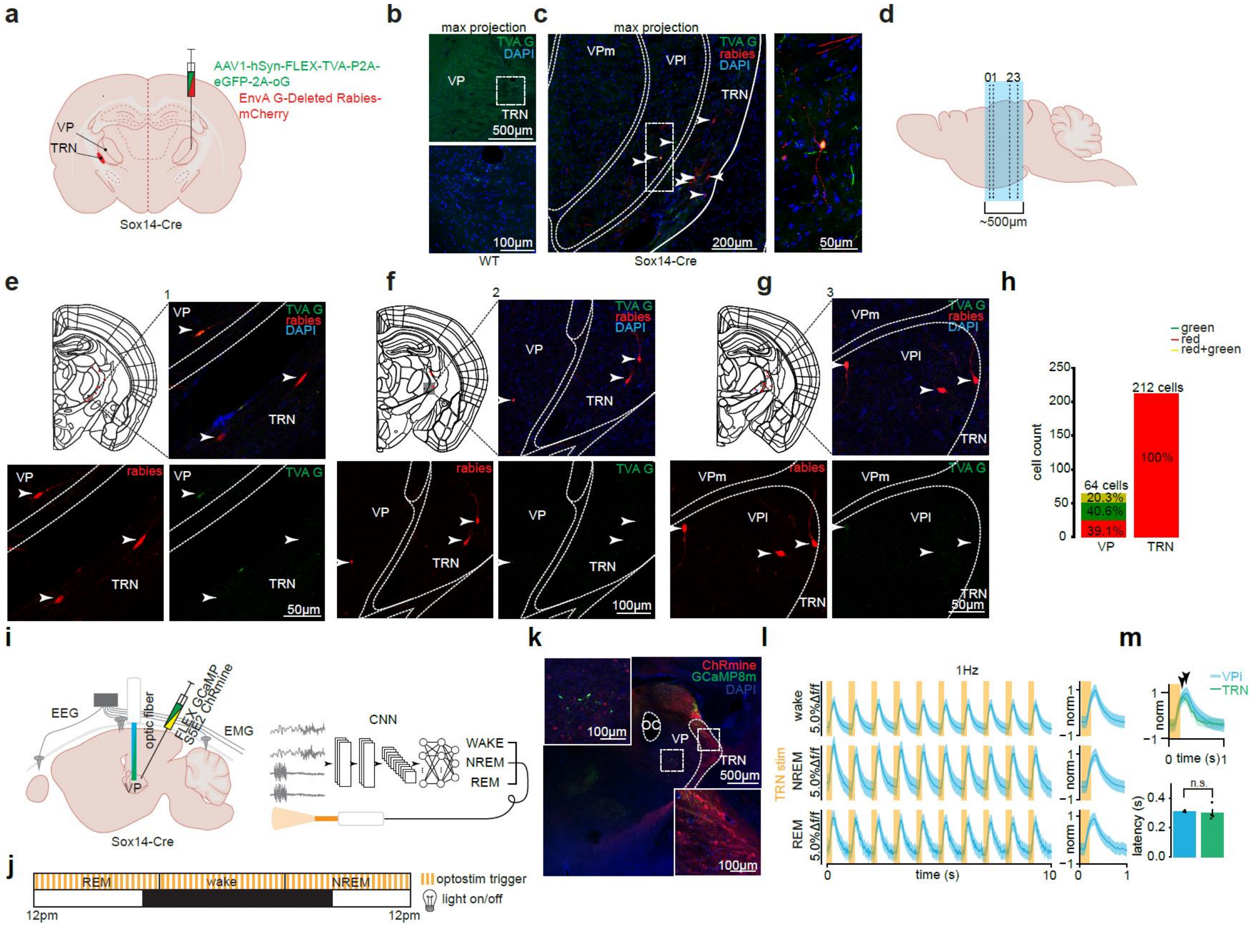


**Extended Data Fig. 8|VPi neurons receive TRN inhibition.**

**a,** Sox14-Cre mice were injected with a helper virus and subsequently injected with rabies into the right hemisphere VP. **b,** Wildtype mice were also injected with the helper virus. No neurons with GFP expression were found. Top: maximum projection image of red and green channels of VP and TRN. Bottom: zoom of inset from top panel. Some green autofluorescence was identified but were too small to be cells and did not have nuclei stained with DAPI (WT control, n = 1 mouse). **c,** Left: max projection image of red, green and blue channels showing the VPl, VPm (ventral posterolateral and ventral posteromedial, subnuclei of the VP), and TRN. Arrows point to all detected neurons. Right: Inset of left panel. A small cell body and one clear VPi morphology. Both neurons have red and green fluorescence (Sox14-Cre, n = 1 mouse). **d,** Lateral view of mouse brain for panels C and E-G. slice "0" is shown in panel C. slices "1-3" are shown in E-G (not to scale). **e-g,** Allen adult mouse brain atlas [68] with neurons located in the VP or TRN. Cells with red fluorescence are shown with red circles, cells with red+green fluorescence are shown with black circles and cells with green fluorescence are shown with blue circles. Grey inset in reference atlas is zoomed in the following 3 associated images. **h,** Cell counts for 15 consecutive slices. **i,** Sox14-Cre mice were injected with a FLEX-GCamp8m and S5E2-ChRmine. **j,** Optostimulation protocols were performed as before (see Fig. 4). **k,** VPi neurons expressed GCaMP8m and TRN neurons expressed ChRmine. OC = optic fiber cannula. **l,** 100 randomly sampled 10s segments were drawn from NREM and wake. 20 samples of REM were taken due to their sparsity. Random sampling iterations occurred 100 times. Segments with mean values exceeding a z-score threshold of 1 were excluded from analysis. Optostimulation of TRN neurons during any behavioral state enabled rebound bursts of VPi neurons. (*n* = 3 mice). **m,** Arrows show the approximate peak location. The response latency of TRN neurons from TRN stimulation at 1Hz (0.2999±0.02469s; TRN dataset is a subset from Fig. 4e) is compared to the response latency of VPi neurons with TRN stimulation (0.3110±0.0041s; n.s. one-sided Mann-Whitney U test). n.s. = not significant.

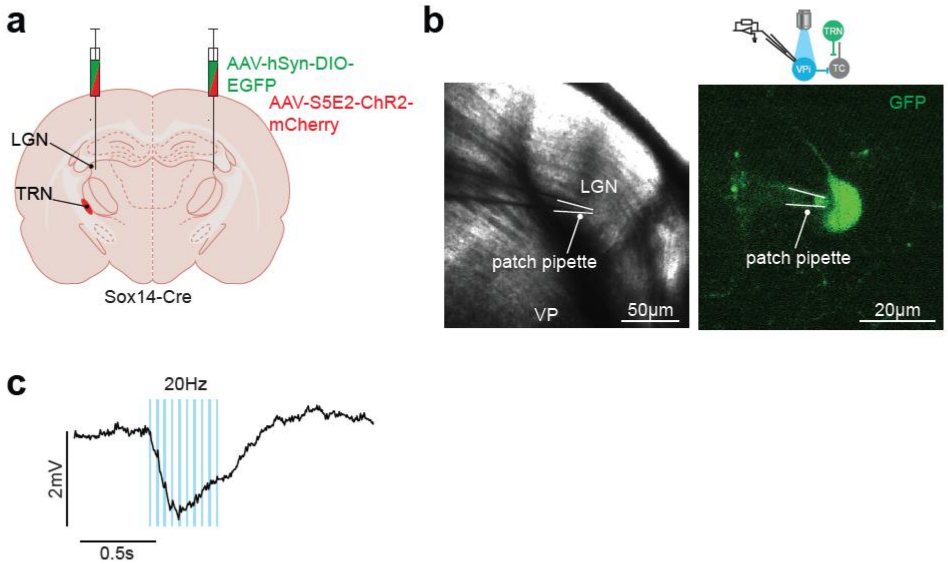


**Extended Data Fig. 9|LGNi neurons receive TRN inhibition.**

**a,** Stereotaxic injection of AAV-hsyn-DIO-EGFP and AAV-S5E2-ChR2-mCherry (bilateral) near the LGN of Sox14-Cre mice. **b,** brightfield and 2 photon image of whole cell patched LGNi. **c,** Averaged LGNi activity during 20Hz blue light stimulation. *n* = 2 neurons.
